# A non-canonical role of *Myc* in programming basal cells as sentinels of upper airway immunity during influenza infection

**DOI:** 10.64898/2025.12.22.696065

**Authors:** Alexander G. Foote, Le Xu, Jamie Verheyden, Belle Pan, Nikita Katoch, Xin Sun

## Abstract

The upper respiratory epithelium is a specialized barrier that integrates immune surveillance with the biomechanical requirements of the laryngotracheal axis. Despite the clinical prevalence of persistent airway dysfunction following viral infection, the region-specific mechanisms governing laryngotracheal injury and repair remain poorly defined. Using integrated mouse models, imaging, and single-cell transcriptomics, we dissected IAV-induced injury in the upper respiratory tract. We found that infection displays regionally restricted tropism to subglottic and tracheal pseudostratified epithelia, targeting ciliated, secretory, neuroendocrine, and basal cells. Rapid viral clearance was accompanied by acute neutrophil infiltration and the emergence of a previously unrecognized intraepithelial CD8^+^ natural killer T (NKT)-like effector population. Functional studies revealed that injury-induced *Myc* expression in KRT5^+^ basal progenitors is required for mucosal restoration; while basal-specific *Myc* deletion preserved viral clearance, it attenuated proliferation and impaired the recruitment of CD8^+^ NKT cells. Together, these findings delineate a coordinated epithelial–immune circuit governing post-viral mucosal restoration, positioning basal cells as specialized epithelial sentinels that orchestrate acute repair and recruit cytotoxic effectors via a MYC-dependent CXCL10–CXCR3 axis.

## Introduction

The upper respiratory epithelium serves as a frontline barrier defense, shielding the airways from a constant barrage of symbiotic microbes, environmental irritants, and a range of invading pathogens.^1-4^ Mucosal immunity is critical to maintain organ homeostasis following epithelial insult.^5,6^ Among common respiratory pathogens, influenza A virus (IAV) remains a major global threat, causing substantial morbidity, acute respiratory distress, and death each year.^7^ While epithelial cells exhibit significant regenerative capacity, it is not well-described how the upper respiratory epithelium remodels in post-viral settings.

The larynx and proximal trachea form a specialized, yet understudied, segment of the upper airway in which epithelial barrier function must be precisely coordinated with biomechanical control amid constant immunologic challenges. Work in papillomaviral disease has highlighted this complexity: in severely immunocompromised mice, laryngeal infection with MmuPV1 leads to severe dysplasia and invasive cancer persisting for months,^8^ whereas immunocompetent mice demonstrate protection against recurrent or vertical infection,^9^ underscoring the importance of epithelial–immune crosstalk in shaping local antiviral responses. Yet, beyond these models, the impact of other clinically relevant viruses—such as influenza—on laryngeal and upper airway biology remain surprisingly unexplored, with limited data on multi-organ, systems-level analysis.

Recent single-cell studies have revealed the upper respiratory epithelium to be highly compartmentalized, with distinct proximal–distal and basal–luminal transcriptional programs across the pharyngolaryngeal and tracheobronchial regions.^10^ Consistent with this organization, emerging airway studies have identified regionally specialized epithelial niches with enhanced regenerative capacity following injury.^11^ Many respiratory viruses enter through, and initially engage with, sentinel epithelial cells at these mucosal surfaces.^6^ Early repair responses trigger inflammatory cascades that are central to local immune regulation and, when dysregulated, contribute to chronic disease.^1,6^ Acute upper respiratory infections are among the most common illnesses globally;^12^ although typically self-limiting, a subset of patients experience persistent sensory and motor dysfunction long after acute illness, suggesting systems-level changes. Post-viral vagal neuropathy exemplifies this phenomenon, often manifesting as chronic cough, throat clearing, dysphonia, vocal fold paresis, and vocal fatigue.^13-18^ Despite strong clinical associations linking these symptoms to antecedent viral illness, the mechanisms driving post-viral dysfunction remain undefined.

To address this gap, we sought to define the upstream cellular and molecular events that shape epithelial repair and immune activation following IAV-induced injury in the mouse upper respiratory tract. Using an integrative approach combining mouse genetics, cell-based assays, and single-cell transcriptomics, we uncovered coordinated epithelial and immune responses that extends well beyond the acute phase of infection. We show that IAV infection targeting discrete epithelial lineages (ciliated, secretory, neuroendocrine, and basal) elicits a rapid Ly6G^+^ neutrophilic response and establishes a persistent intraepithelial population of cytotoxic CD8^+^ NKT-like cells marked by a durable effector transcriptome. We further identify ectopic *Myc* induction within injured subglottic and tracheal pseudostratified epithelium and demonstrate that KRT5^+^ basal progenitors are programmed as epithelial sentinels that deploy a *Myc*-dependent proliferative and chemokine response to shape downstream immune behavior. Basal-specific loss of *Myc* blunts epithelial expansion and disrupts lineage differentiation, while attenuating CXCL10-dependent recruitment of CXCR3^+^ CD8^+^ NKT cells. Together, these findings delineate a systems-level epithelial– immune circuit in which basal cells act as sentinels of upper airway immunity during influenza infection and offer mechanistic insight into clinically relevant sequelae such as post-viral vagal dysfunction.

## Results

### Spatiotemporal dynamics of IAV infection reveal regionally restricted epithelial tropism and rapid viral clearance

To delineate the spatiotemporal progression of IAV infection along the upper respiratory tract, we generated encompassing large-scale the coronal sections pharyngolaryngeal-to-tracheobronchial axis from wild-type (WT) mice at 1, 3, 7, and 8 days post-infection (DP-IAV) (Fig. 1a,b). Multiplex RNA in situ hybridization with probe targeting IAV transcripts revealed initial viral localization to the airway epithelium at 1DP-IAV, with peak epithelial coverage at 3DP-IAV (Fig. 1c,d). Viral transcripts persisted through 7DP-IAV but were eradicated by 8DP-IAV, aligning with established timelines of innate immune-mediated clearance.^19^ Consistent with prior reports,^20^ IAV induced transient morbidity with decrease in body weight of ∼20%, reaching its lowest point at 10DP-IAV before returning to near baseline levels by 21DP-IAV (Fig. 1e).

**Fig. 1.**
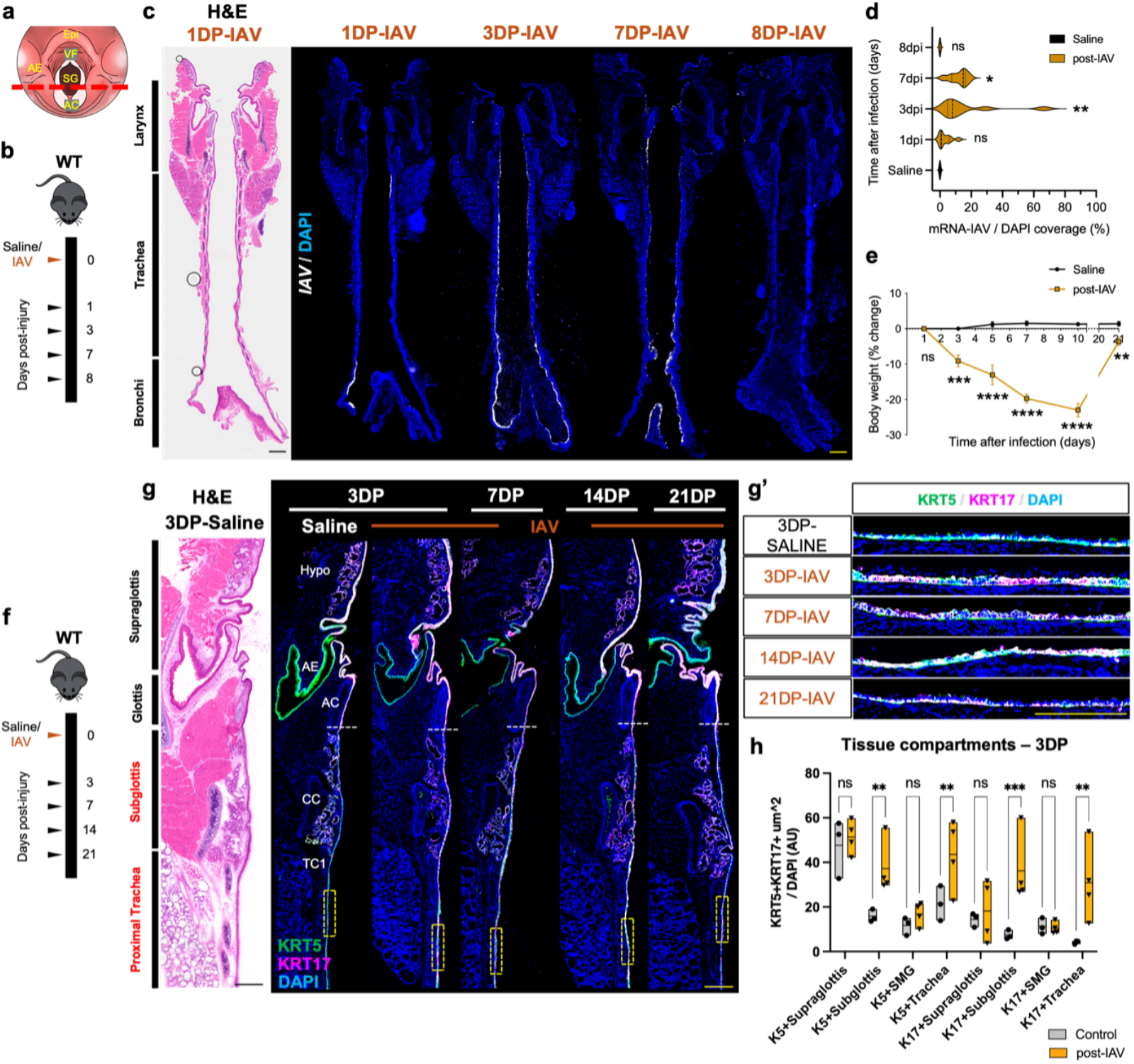
IAV infection reveals regionally restricted epithelial injury and KRT5^+^ basal cell–mediated repair. **a**,**b**, Schematics of upper airway coronal sectioning and IAV infection timeline. **c**, H&E and RNAscope detection of IAV-mRNA (white) at 1, 3, 7, and 8DP-IAV. **d**, Quantification of epithelial IAV-mRNA coverage (one-way ANOVA with Dunnett’s post-hoc test; mean ± SEM; n = 3/group). **e**, Body weight change following infection (Welch’s t-test; mean ± SEM; n = 8/group). **f**, Timeline schematic corresponding to panels g–g′. **g-g′**, H&E and immunofluorescence for KRT5 (green) and KRT17 (magenta) showing basal cell hyperplasia at 3–14DP-IAV with resolution by 21DP-IAV; dashed boxes indicate higher-magnification insets. **h**, Quantification of KRT5/KRT17 epithelial coverage across airway compartments (two-way ANOVA with Fisher’s LSD post-hoc test; mean ± SEM). DAPI, blue. Scale bars: 500 µm (c,g) 100 µm (others). *P < 0.05, **P < 0.005, ***P < 0.0005; ns, not significant; WT, wild-type; DP, days post.

Building on this temporal framework, we interrogated regional vulnerabilities by reorienting sections along dorsal-ventral and medial-lateral axes at peak infection (3DP-IAV). Pseudostratified epithelia of the subglottis, trachea, and extrapulmonary bronchi exhibited the highest susceptibility to IAV-induced cytopathy, whereas stratified, squamous regions of the glottis and supraglottis harbored fewer infected cells (Fig. 1c; Fig. S1a). This topographic bias suggested that epithelial architecture and cell-type composition dictate viral entry and propagation.

To link infection dynamics to regenerative potential, we traced basal progenitor responses using immunofluorescence for keratins KRT5 and KRT17—hallmarks of proliferative basal cells—across acute (3–7DP), subacute (14DP), and resolution (21DP) phases (Fig. 1f). Hyperplasia of KRT5^+^KRT17^+^ cells emerged by 3DP-IAV, crested at 7–14DP-IAV, and normalized by 21DP-IAV (Fig. 1g,g′). Compartment-specific quantification (supraglottic surface epithelium [SE], subglottic SE, tracheal SE, and submucosal glands [SMG]) confirmed that expansion was confined to pseudostratified zones of prior viral enrichment, with maximal accrual in subglottic and tracheal epithelia (Fig. 1h; Fig. S1b). Thus, basal cell activation represents a targeted reparative program, spatially attuned to sites of epithelial compromise.

We next resolved the cellular substrates of infection through colocalization of IAV transcripts with lineage-defining markers (Fig. 2). In the trachea, viral RNA predominantly colocalized with FOXJ1^+^ ciliated and SCGB1A1^+^ club cells (Fig. 2c), while in the laryngeal epiglottis, SCGB1A1^+^ secretory cells were preferentially targeted (Fig. 2d). Lineage tracing in Ascl1^CreER;^tdTomato mice further implicated subglottic and tracheal neuroendocrine cells as viral reservoirs (Fig. 2e). Additionally, IAV enrichment extended to KRT13− suprabasal layers of stratified squamous hillocks, arytenoid, and vocal fold mucosa (Fig. 2f-h), as well as dispersed KRT5^+^Ki67^+^ basal cells (Fig. 2i). Collectively, these data delineate a broad yet selective epithelial tropism, wherein IAV exploits regionally specialized sentinel cells to establish foothold, culminating in viral detection and progenitor-driven remodeling.

**Fig. 2.**
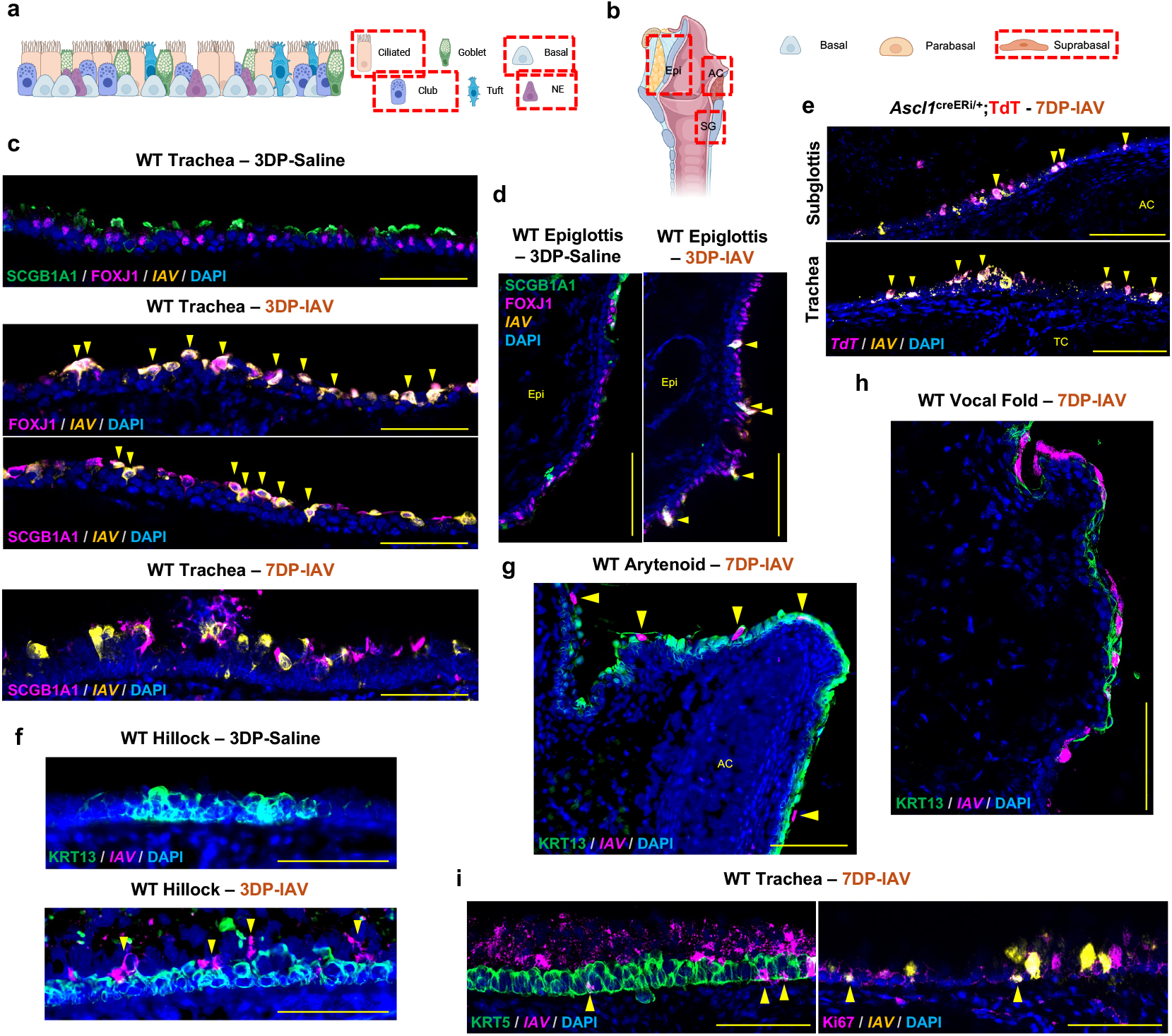
IAV demonstrates broad cellular tropism while encountering restricted entry into stratified squamous epithelium. **a**,**b**, Schematics of airway epithelium lineages and laryngeal anatomy, delineating pseudostratified zones from stratified squamous epithelial (SSE) compartments. **c**, IF of WT trachea at 3 and 7DP-IAV; IAV^+^ signal (arrowheads) predominates in FOXJ1^+^ ciliated and SCGB1A1^+^ club cells. **d**, Comparative analysis of pseudostratified epithelial regions of the WT epiglottis at 3DP-IAV revealing similar lineage-restricted infection dynamics. **e**, Ascl1^creER/+^;TdT lineage tracing confirms IAV tropism for subglottic and tracheal neuroendocrine cells. **f-h**, IAV infection in stratified squamous epithelia of the tracheal hillock, laryngeal arytenoid, and vocal fold is confined to KRT13^−^ suprabasal cells. **i**, Co-staining for IAV and KRT5 or Ki67 in the trachea identifies a subset of infected basal progenitors during peak injury. DAPI, blue. Scale bars: 100 µm. WT, wildtype; DP, days post; epiglottis, Epi; arytenoid cartilage, AC; subglottis, SG; tracheal cartilage, TC.

### Acute neutrophil infiltration and the emergence of a persistent CD8^+^ NKT-like population characterize the post-IAV laryngotracheal niche

Given that viral clearance hinges on coordinated innate and adaptive immunity to preserve epithelial barrier function,^5^ we systematically profiled immune infiltration to dissect its contributions to resolution and homeostasis. Prior studies have implicated neutrophils in early antiviral containment and CD8^+^ T cells in late-stage cytotoxicity;^21,22^ thus, we prioritized these compartments using Ly6G (neutrophil) and CD3 (pan-T cell) immunostaining in WT tissues at acute (3DP-IAV, 7DP-IAV) and chronic (21DP-IAV) stages (Fig. 3a–f). In uninfected controls, Ly6G^+^ neutrophils resided in the subepithelial compartment. At 3DP-IAV, Ly6G^+^ neutrophils showed massive infiltration into the subglottic and tracheal epithelia, colocalized with IAV enriched cells, forming dense aggregates within the luminal space (Fig. 3d,d′; Fig. S1c). This acute influx coincided with maximal epithelial denudation and peak viral burden and resolved by 7DP-IAV (Fig. 3d′), supporting a model in which neutrophil epithelial invasion is confined to early antiviral containment.

**Fig. 3.**
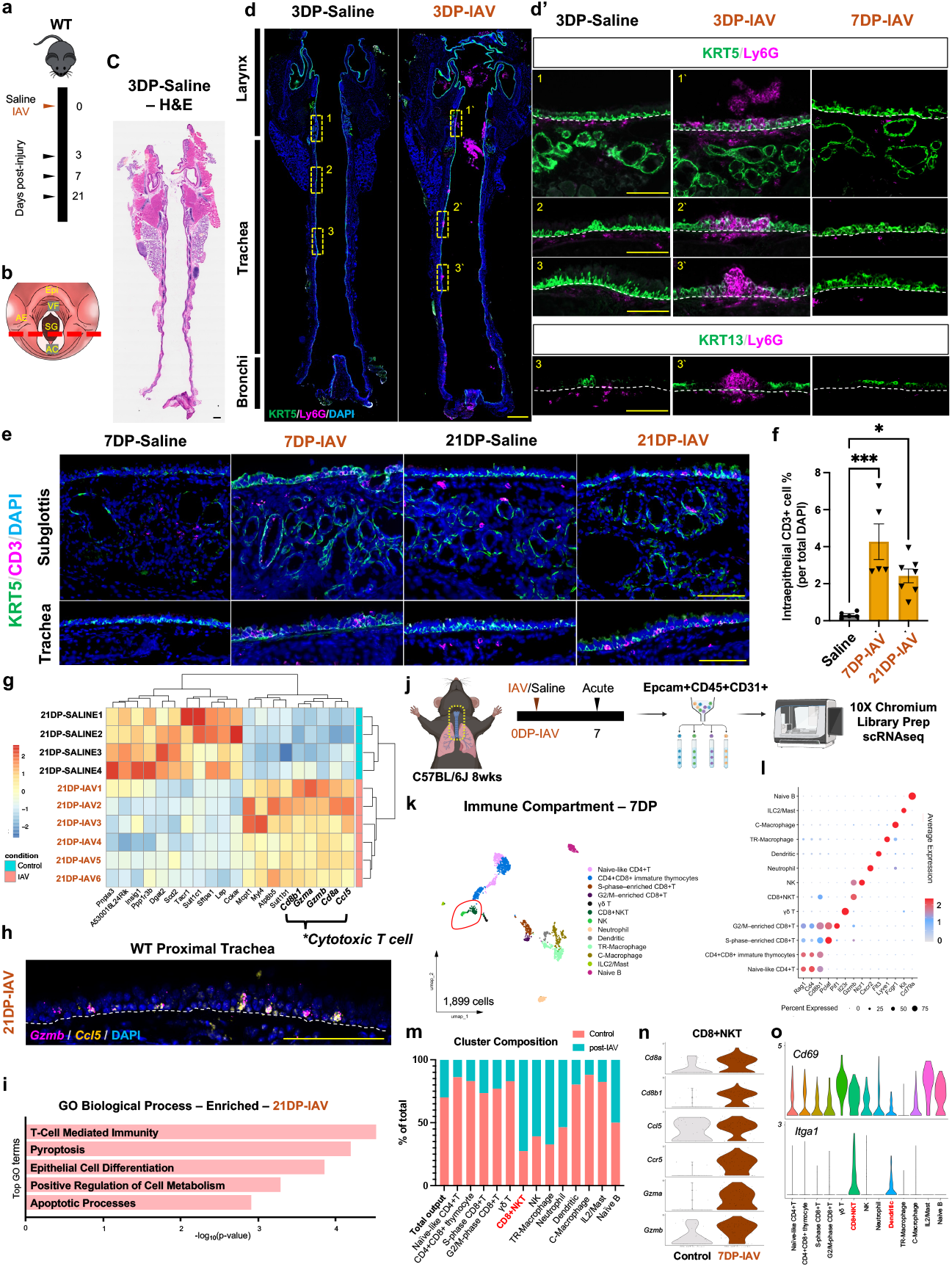
Acute IAV infection recruits neutrophils and establishes persistent intraepithelial CD8^+^ NKT cells. **a**,**b**, Schematics of IAV infection timeline and airway sectioning. **c**, H&E staining of sa-line-treated controls at 3 DP. **d**,**d′**, Immunofluorescence for Ly6G^+^ neutrophils at 3 and 7 DP-IAV demonstrating epithelial infiltration; dashed boxes indicate insets. **e**, Immunofluorescence showing CD3^+^ intraepithelial T cells in subglottic and tracheal epithelium at 7 DP-IAV that persist at 21 DP-IAV. **f**, Quantification of CD3^+^ epithelial coverage (one-way ANOVA with Dunnett’s post-hoc test; mean ± SEM). **g**, Bulk RNA-seq showing increased expression of cytotoxic T-cell genes (Cd8a, Cd8b1, Ccl5, Gzma, Gzmb) at 21 DP-IAV. **h**, RNAscope co-localization of Gzmb and Ccl5 transcripts at 21 DP-IAV. **i**, Top enriched GO biological processes from bulk RNA-seq at 21 DP-IAV. **j**, Schematic of scRNA-seq workflow. **k**, UMAP of immune populations at 7 DP-IAV (n = 8 biological replicates). **l**, Dot plot of top cluster markers. **m**, Immune cluster composition comparing saline and IAV. **n**,**o**, Violin plots showing enrichment of cytotoxic and tissue-resident gene signatures in CD8^+^ NKT cells. DAPI, blue. Scale bars: 500 µm (c,d) 100 µm (others). *P < 0.05, **P < 0.005, ***P < 0.0005; ns, not significant; WT, wild-type; DP, days post.

In contrast, intraepithelial CD3^+^ T cells accrued progressively, peaking at 7DP-IAV and remaining significantly expanded at 21DP-IAV over saline controls (Fig. 3e,f). To molecularly characterize this resident population, bulk RNA sequencing of 21DP-IAV laryngeal and tracheal tissues identified a cytotoxic signature dominated by Cd8a, Cd8b1, Gzma, Gzmb, and Ccl5 transcripts (Fig. 3g). In situ validation confirmed colocalization of Gzmb+Ccl5^+^ cells within the epithelium at 21DP-IAV (Fig. 3h). Gene Ontology (GO) analysis of differentially upregulated genes was enriched for T cell-mediated immunity (Cd8a, Prf1), pyroptosis (Gzma, Gzmb), epithelial differentiation (Sult1b1, Dhrs9, Cdk1), metabolic regulation (Ccl5, Cdk1), and apoptosis (Gzma, Prf1, Gzmb) (Fig. 3i), implying multifaceted contributions to both viral lysis and tissue restitution.

To integrate these immune dynamics with epithelial repair and viral hotspots, we conducted single-cell RNA sequencing (scRNA-seq) on en bloc-dissociated tissues along the pharyngolaryngeal-to-tracheobronchial axis from 7DP-IAV—the timepoint between peak inflammatory repair and viral clearance (Fig. 3j). Unbiased clustering resolved 13 immune subsets (Fig. 3k,l; Fig. S1d-g), with compositional shifts post-IAV dominated by T/NK lineage expansion (Fig. 3m). Notably, we identified a transcriptionally distinct CD8^+^ natural killer T (NKT)-like cell cluster, co-expressing canonical T cell (Cd3d, Cd8b1), NK (Nkg7), and Th1/iNKT markers (Klrk1, Tbx21, Ifng), alongside cytotoxic effectors (Gzma, Gzmb, Ccl5) (Fig. 3l,n; Fig. S1g). Consistent with a tissue-resident phenotype, these cells upregulated canonical residency markers (Cd69, Itga1) and mirrored transcriptional programs observed in our bulk-seq dataset (Fig. 3n,o), suggesting that CD8^+^ NKT cells may function as durable effectors for immune surveillance.

### scRNA-seq uncovers MYC-driven proliferative basal progenitors in post-IAV epithelial repair

To dissect the epithelial regenerative repertoire during the transitional phase of IAV clearance, we interrogated EpCAM^+^-enriched fractions from our 7DP scRNA-seq atlas (Fig. 4a). Dimensionality reduction and unsupervised clustering delineated 20 distinct transcriptional states spanning the upper respiratory epithelium (Fig. 4a–d; Fig. S1h), which segregated into laryngeal, tracheal, and SMG domains in accordance with anatomical and prior transcriptomic delineations.^10^ A salient feature was the emergence of proliferative epithelial clusters, disproportionately represented in our post-IAV condition (Fig. 4b,c,e), implying a stimulus-dependent expansion or transdifferentiation program in response to viral perturbation.

**Fig. 4.**
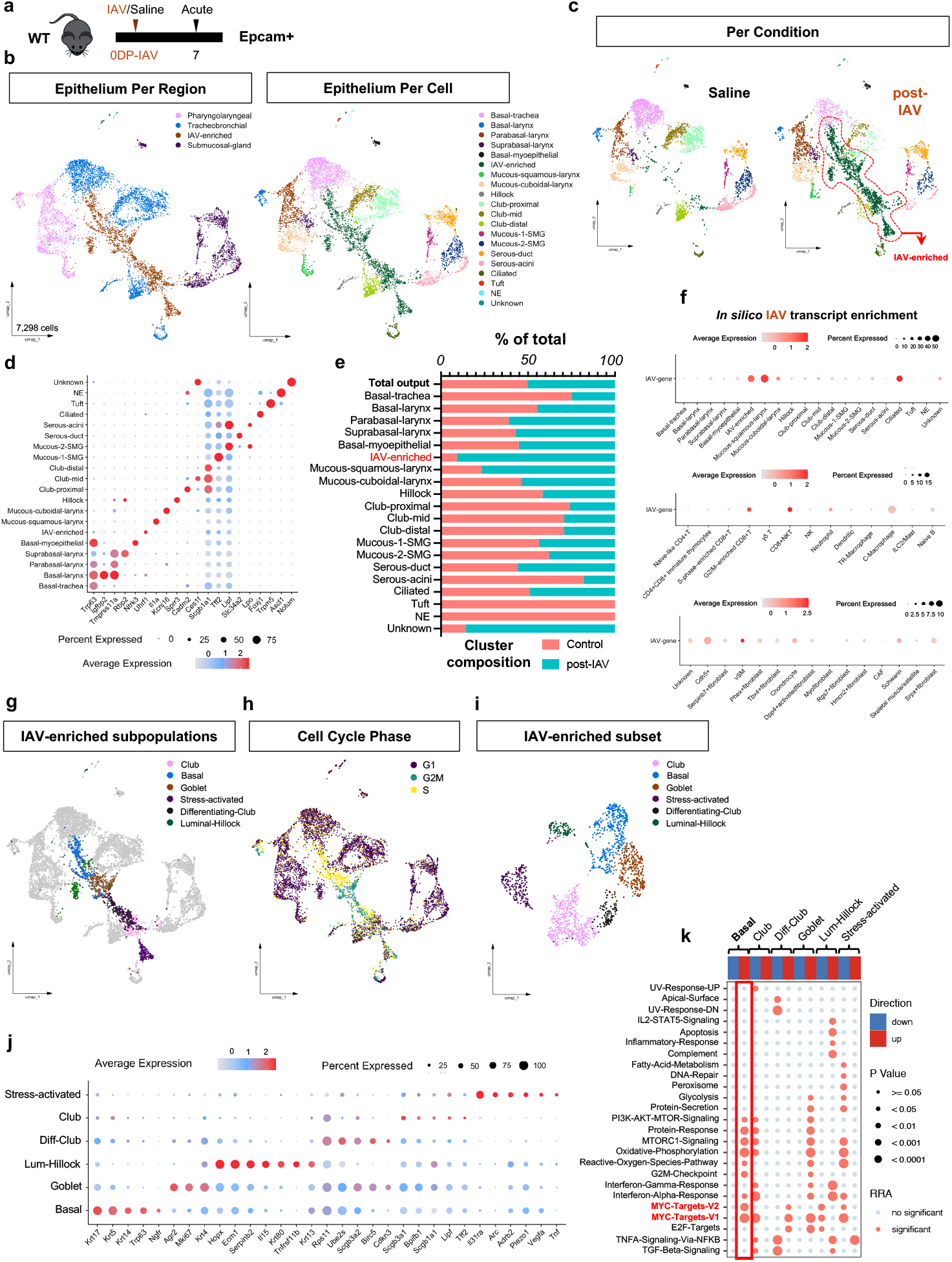
scRNA-seq identifies *Myc*-driven basal progenitor programs within IAV-enriched epithelial populations. **a**, Schematic of scRNA-seq experimental design. **b**, Integrated UMAP of upper airway epithelial populations highlighting regional and cell-type diversity. **c**, UMAP colored by condition (saline vs IAV) identifying IAV-enriched epithelial clusters. **d**, Dot plot of top markers defining epithelial clusters. **e**, Cluster composition comparing saline and IAV conditions. **f**, *In silico* viral transcript mapping demonstrating preferential IAV enrichment in epithelial lineages at 7DP-IAV. **g-i**, UMAPs highlighting IAV-enriched epithelial subsets and associated cell-cycle states (G1, S, G2M). **j**, Dot plot of markers defining IAV-enriched clusters. **k**, Integrated regulatory GSEA identifying activation of MYC target pathways in IAV-enriched basal progenitors. WT, wild-type; DP, days post.

To pinpoint viral niches at cellular resolution, we augmented the reference transcriptome with the complete IAV genome sequence (TableS1), enabling imputation of viral transcripts across clusters. This revealed a stark epithelial tropism, with highest IAV-mRNA enrichment in secretory, ciliated, and—critically—the novel IAV-enriched (e.g. responsive) subsets (Fig. 4f; TableS1), thereby validating our spatial mapping. Sporadic detection in CD8^+^ NKT cells (<15%) and vascular smooth muscle (<5%) hinted at intercellular viral transfer or bystander activation (Fig. 4f).

To exclude the possibility that incorporation of viral transcripts biased dimensionality reduction or clustering, we reprocessed the scRNA-seq dataset after removing the synthetic IAV reference while holding all parameters constant. Cell-type identities and manifold structure were fully preserved, with identical clustering outcomes and near-perfect concordance across embeddings (Fig. S2a; Table S1). These analyses confirm that the emergence of IAV-enriched epithelial states reflects biological responses to infection rather than computational artifacts introduced by viral transcript mapping.

The temporal coincidence of our IAV-enriched subpopulation with peak basal hyperplasia prompted us to posit a progenitor-like identity underpinning repair. Subclustering unmasked proliferative heterogeneity, with G2/M- and S-phase signatures evoking cell-cycle traversal (Fig. 4g–i; TableS1). Preeminent among these was a basal-enriched contingent, defined by Trp63, Krt5, Krt14, Krt17, and Ngfr, and biased toward S-phase progression—hallmarks of replicative competence in response to genotoxic stress (Fig. 4g-j). Pathway analysis through irGSEA unveiled marked upregulation of MYC target gene sets in the IAV-enriched basal subpopulation (Fig. 4k), inspiring targeted follow-up experiments.

### Basal-specific Myc ablation impairs proliferative repair and lineage commitment following IAV injury

The proto-oncogene transcription factor Myc constitutes a central regulator of immune cell proliferation, metabolic reprogramming, and differentiation, coordinating rapid cellular responses during activation and inflammatory states.^23-25^ Interrogation of our scRNA-seq dataset revealed selective enrichment of Myc—but not L-Myc or N-Myc—within the IAV-enriched population, with the highest expression in the basal subpopulation, where 72% of basal cells expressed Myc (Fig. 5a,b; Fig. S2b,c). Pseudotemporal ordering further supported a stem-like trajectory, bifurcating toward goblet, luminal-hillock, differentiating club, mature club, and stress-activated epithelial fates (Fig. 5c). Spatial validation via in situ hybridization confirmed widespread ectopic Myc expression throughout the pseudostratified epithelium spanning the subglottis, trachea, and extrapulmonary bronchi during the acute post-IAV phase (Fig. 5d). Collectively, these findings designate a MYC-driven basal cell program as a central mediator of acute epithelial regeneration following IAV injury. To assess the functional requirement of Myc in epithelial repair, we generated tamoxifen-inducible basal cell–specific knockout mice (Krt5^creER^;Myc^flox/flox^, hereafter Krt5^creER^;Myc). Experimental animals received three consecutive tamoxifen doses five days prior to IAV inoculation, with tissue harvest performed at 7 and 21DP-IAV (Fig. 5e). Immunofluorescence microscopy with quantification of Myc-mRNA coverage confirmed efficient epithelial Myc deletion (Fig. 5f,g). Notably, Myc deletion in KRT5^+^ basal cells did not alter laryngotracheal epithelial architecture or cellular composition under homeostatic conditions (Fig. S2d), suggesting that basal-specific Myc is dispensable for steady-state maintenance of the upper airway. Phenotypically, Krt5^creER^;Myc mutants mirrored heterozygous controls in acute morbidity, incurring ∼20% weight loss at 7DP-IAV (Fig. S2e).

**Fig. 5.**
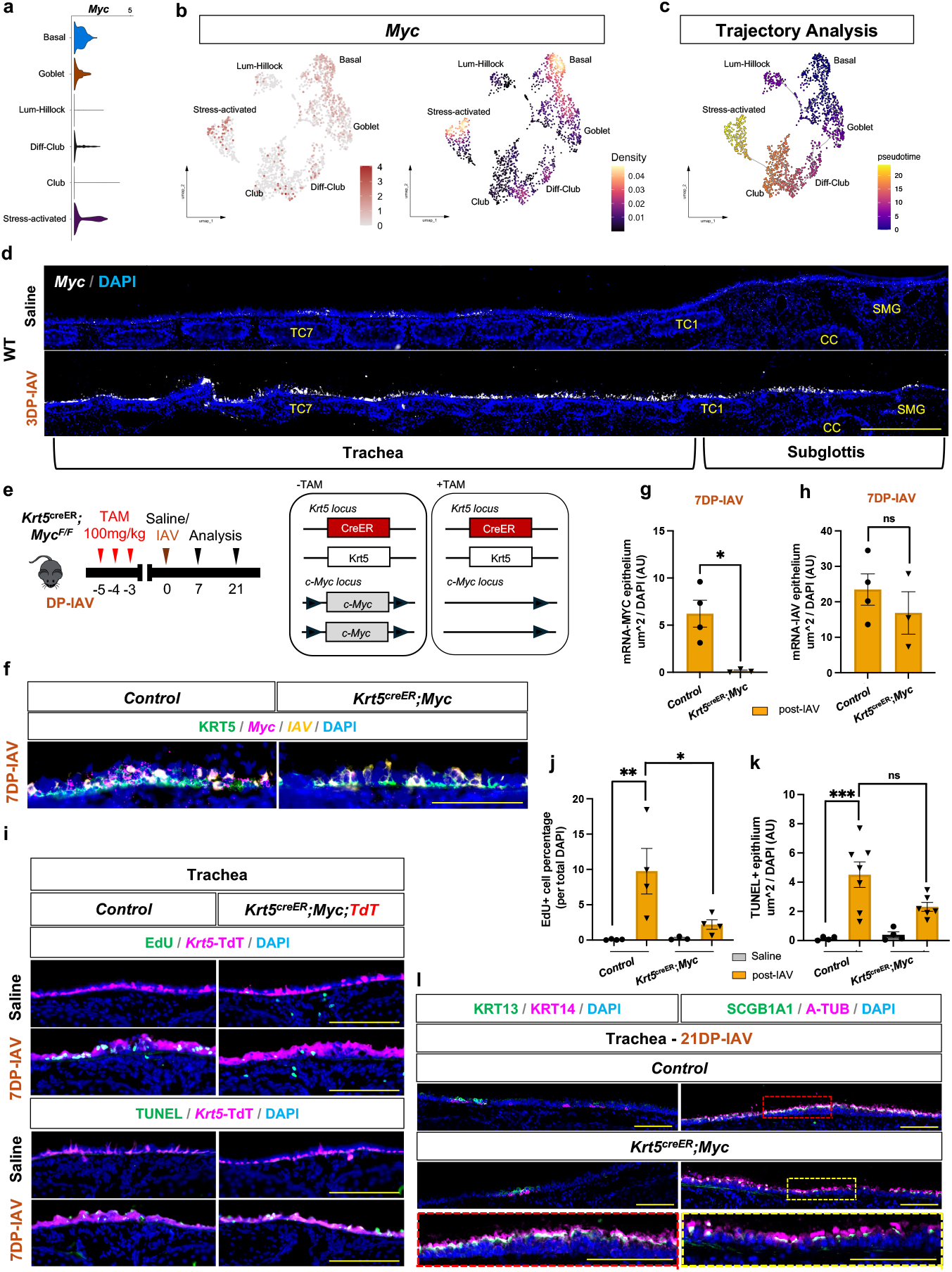
MYC promotes basal progenitor expansion and lineage remodeling during post-IAV epithelial repair. **a**,**b**, Violin and feature plots showing *Myc* expression across IAV-enriched epithelial populations at 7DP-IAV. **c**, Monocle3 pseudotime trajectory analysis of IAV-enriched epithelial subsets. **d**, Immunofluorescence at 3DP-IAV demonstrating ectopic *Myc* expression in damaged pseudostratified epithelium. **e**, Experimental schematic for basal cell–specific *Myc* deletion. **f**, Immunofluorescence confirming *Myc* loss following tamoxifen induction. **g**,**h**, Quantification of epithelial *Myc*-mRNA and *IAV*-mRNA coverage (Student’s t-test; mean ± SEM). **i**, EdU and TUNEL staining assessing proliferation and apoptosis at 7DP-IAV. **j**,**k**, Quantification of proliferative and apoptotic indices (one-way ANOVA with Tukey’s post-hoc test; mean ± SEM). **l**, Immunofluorescence of lineage marker–positive cells in tracheal epithelium at 21DP-IAV. DAPI, blue. Scale bars: 500 µm (**d)** 100 µm (others). *P < 0.05, **P < 0.005, ***P < 0.0005; ns, not significant.

To determine whether Myc deletion impacted viral replication and/or clearance, we assessed IAV transcript localization during the reparative phase. At 7DP-IAV, both control and Krt5^creER^;Myc mutants exhibited similar levels of epithelial IAV-mRNA enrichment, and by 10DP-IAV, viral transcripts were undetectable in both groups, indicating normal viral replication and clearance (Fig. 5h; Fig. S2f).

Given the role of Myc in cell cycle regulation,^24,26^ we next evaluated epithelial proliferation and apoptosis using EdU incorporation and TUNEL staining at 7DP-IAV. Homeostatic epithelia showed no differences in EdU or TUNEL positivity between control and mutant mice (Fig. 5i-k). However, following IAV injury, Krt5^creER^;Myc mutants exhibited a marked reduction in epithelial cell proliferation compared to infected controls, while apoptosis levels remained unchanged (Fig. 5i-k). These findings indicate that Myc drives mitogenic surge upon injury, decoupling it from homeostatic turnover, apoptosis, or antiviral orchestration.

Anticipating chronic sequelae from blunted epithelial expansion, we scrutinized 21DP-IAV histology, yet discerned similar cell abundance and no overt dysmorphia in mutants versus uninjured controls (Fig. S3a). Given that all upper airway epithelial lineages originate from KRT5^+^ progenitors (Fig. S3b), we next examined lineage-defined cell types in Krt5^creER^;Myc mutants compared to infected controls to assess sustained architectural repair at 21DP-IAV. In mutant epithelium, we observed trends toward reduced KRT14^+^ basal cells, modest expansion of KRT13^+^ hillock cells, and a decrease in acetylated-tubulin^+^ ciliated cells, although none of these changes reached statistical significance. Notably, we identified a significant reduction in SCGB1A1^+^ club cells in injured mutants, accompanied by disrupted ciliated and club cell distribution at 21DP-IAV (Fig. 5l; Fig. S3c). Collectively, our data indicate that Myc acts as a critical injury-induced regulator of basal progenitor function, coordinating both epithelial expansion and lineage specification. Loss of Myc blunts proliferation and disrupts differentiation trajectories, resulting in mispatterned epithelial remodeling at 21DP-IAV.

### Epithelial-Immune Crosstalk Establishes a MYC-Dependent Cxcl10–Cxcr3 Axis Driving CD8^+^ NKT-like Cell Recruitment Post-IAV

Epithelial cells, as primary targets of IAV-mediated cytopathy, orchestrate immune recruitment via secreted mediators to expedite viral clearance and mucosal restoration.^5,6^ To test the hypothesis that Myc expression in basal cells modulates immune recruitment, we interrogated differentially expressed genes in our scRNA-seq dataset, focusing on IAV-enriched epithelial cells. Cxcl10 emerged as the singular chemokine significantly upregulated, with Myc^+^ co-expression confined to the injury-responsive basal population (Fig. 6a; Fig. S3d), positioning it as a candidate regulator of epithelial-driven immune influx. Consistent with this single-cell signature, Cxcl10 expression in situ closely mirrored viral burden across the upper airway—peaking during active infection and declining to negligible levels by 8DP-IAV—underscoring its role in recruiting effector cells to sites of ongoing viral replication (Fig. S3e).

**Fig. 6.**
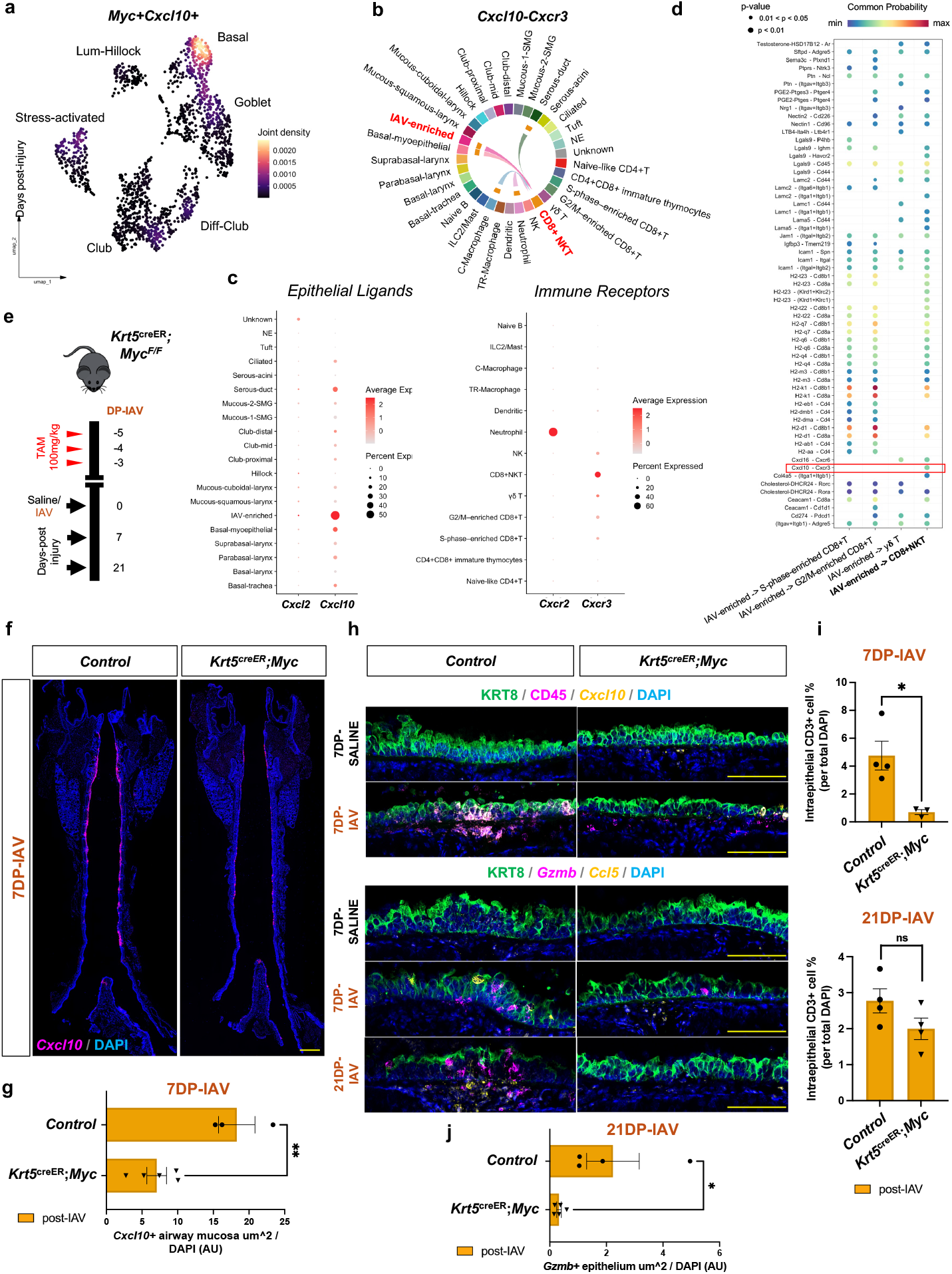
MYC-dependent epithelial CXCL10 signaling recruits CD8^+^ NKT cells during post-IAV repair. **a**, UMAP of IAV-enriched epithelial populations showing co-expression of *Myc* and *Cxcl10* at 7DP-IAV. **b-d**, CellChat analysis identifying CXCL10–CXCR3 as the dominant IAV-enriched-to-CD8^+^ NKT signaling axis. **e**, Schematic of tamoxifen induction and IAV infection timeline in Krt5^CreER^;Myc mutant mice. **f**,**g**, Whole-mount RNAscope imaging and quantification showing reduced epithelial *Cxcl10* expression in Krt5^CreER^;Myc mutants at 7DP-IAV. **h**, Immunofluorescence showing reduced CD45^+^ immune infiltration and epithelial *Gzmb*/*Ccl5* expression in mutants at 7 and 21DP-IAV. **i**, Quantification of intraepithelial CD3^+^ cells. **j**, Quantification of epithelial *Gzmb* expression at 21DP-IAV. DAPI, blue. Scale bars: 500 µm **(f)**, 100 µm (others). *P < 0.05, **P < 0.005, ***P < 0.0005; ns, not significant.

To map epithelial-immune communication, we applied CellChat to infer ligand-receptor interactions.^27^ Cxcl10 transcripts were detected across epithelial (serous-ductal, IAV-enriched, basal-myoepithelial) and immune (TR-macrophage, neutrophil) populations, yet its receptor, Cxcr3, was exclusively expressed in our CD8^+^ NKT cells (Fig. 6b-d). Probabilistic signaling analysis underscored the Cxcl10–Cxcr3 axis as the predominant conduit linking IAV-enriched epithelia to CD8^+^ NKT cells (Fig. 6c,d), implicating this pathway in precise immune recruitment during repair. In contrast, interrogation of the Cxcl2–Cxcr2 axis—previously tied to neutrophil dynamics^22^—revealed no epithelial Cxcl2 expression at 7DP-IAV, excluding its role in epithelial-driven responses at this stage (Fig. 6c).

To interrogate the functional control exerted by Myc over this epithelial–immune chemokine axis, we profiled immune recruitment dynamics in Krt5^creER^;Myc mutants versus heterozygous controls at acute 7DP-IAV and convalescent 21DP-IAV phases (Fig. 6e). In situ hybridization disclosed a profound depletion of Cxcl10-mRNA across the pseudostratified airway mucosa in mutants relative to controls specifically at the acute 7DP-IAV stage (Fig. 6f–h), underscoring the requisite role of Myc in injury-induced chemokine induction. This blunted CXCL10 gradient translated to impaired mucosal immunity whereby mutants displayed significantly reduced CD45^+^ leukocyte and CD3^+^ T-cell infiltration at 7DP-IAV (Fig. 6h,i; Fig. S3f), with trends toward persistence at 21DP-IAV that fell short of statistical significance. Although overall CD3^+^ accrual normalized by 21DP-IAV, epithelial expression of hallmark CD8^+^ NKT effector genes—Gzmb and Ccl5—was durably subdued in mutants. Notably, Gzmb transcripts exhibited sustained, statistically significant attenuation at 21DP-IAV (Fig. 6h,j), implying that Myc orchestrates not only initial recruitment but also the long-term priming and cytotoxic arming of intraepithelial effectors. Collectively, these findings establish that Myc orchestrates a Cxcl10–Cxcr3– dependent epithelial–immune signaling circuit, enabling selective recruitment and functional maturation of CD8^+^ NKT cells for coordinated mucosal repair following IAV injury.

## Discussion

Viral infections in the airway play a pivotal role in modulating host immune responses, influencing both the resolution of infection and the preservation of epithelial integrity essential for airway health and function. Here, we define a MYC-dependent epithelial–immune circuit that orchestrates repair of the upper respiratory tract after IAV injury. Using spatial mapping, scRNA-seq, and conditional genetics, we show that IAV targets regionally specialized epithelial lineages and elicits a MYC-driven basal progenitor response that is coupled to recruitment of cytotoxic CD8^+^ NKT effector cells via a CXCL10–CXCR3 axis. Basal cell–specific *Myc* deletion uncouples these processes such that viral replication/clearance and gross architecture are preserved, but injury-induced proliferation, lineage allocation, and cytotoxic T cell recruitment are attenuated, resulting in subtle yet mispatterned remodeling of secretory and ciliated compartments.

Our findings extend a growing literature positioning *Myc* as a central regulator of epithelial regeneration beyond its canonical oncogenic role.^25,26,28,29^ *Myc* has been implicated in coordinating proliferation and metabolic rewiring in diverse regenerative contexts, including intestinal, hepatic, and skeletal tissues.^30-32^ In the lung, *Myc* promotes activation of the peribronchiolar smooth muscle cell niche and *Fgf10* expression after naphthalene injury, and its deletion in this compartment attenuates epithelial repair.^33^ Work in corneal epithelium further demonstrates *Myc* maintains epithelial architecture at homeostasis and governs the balance among stratification, differentiation, and exfoliation during wound repair, in part through modulation of *p63*-dependent cell-fate programs.^26^ Our data add a complementary layer by showing that, in the upper respiratory tract, *Myc* is selectively induced in KRT5^+^ basal progenitors of laryngotracheal pseudostratified epithelia after viral injury and is required for a full proliferative burst and appropriate differentiation into club and ciliated lineages. Across epithelial organs, *Myc* thus emerges as a conserved coordinator of injury-induced regeneration whose precise regulation is essential for maintaining tissue homeostasis. Whether this program is conserved across non-viral airway challenges such as chemical/environmental irritants and bacterial infection remains to be determined. In homeostatic conditions, basal cells of the stratified squamous laryngeal epithelium displayed robust *Myc* expression; nonetheless, this region experienced negligible viral insult, precluding any detectable MYC-mediated repair program. Accordingly, we propose that similar MYC-regulated programs exist in this niche but are engaged only after substantial epithelial compromise.

A key feature of our model is the integration of the MYC-driven basal progenitor program with localized cytotoxic immunity. We observe an intraepithelial CD3^+^ T cell population persisting at 21DP-IAV that displays a tissue-resident, CD8^+^ NKT–like cytotoxic transcriptional signature not previously described in the upper airway. scRNA-seq and CellChat analysis revealed that *Myc*+ basal and IAV-enriched epithelial cells are major sources of the chemokine *Cxcl10*, whereas its receptor *Cxcr3* is largely confined to these CD8^+^ NKT cells, positioning this axis as a dominant conduit of epithelial–immune crosstalk. The CXCL10–CXCR3 axis is well recognized for directing effector T-cell trafficking to injured tissues.^34-39^ In influenza models, CXCR3^+^ CD8^+^ T cells and lung tissue-resident memory (T_RM_) cells provide rapid heterosubtypic protection upon re-exposure, yet subsets of CXCR3^+^ or T_RM_ CD8^+^ T cells can also exacerbate chronic lung pathology without significantly altering viral clearance, indicating that functional outcomes are context dependent.^36^ Consistent with this complexity, *Myc*-deficient airways show reduced epithelial *Cxcl10* expression and diminished early CD3^+^ recruitment, indicating that MYC tunes both the magnitude and positioning of cytotoxic cells during repair.

Several caveats warrant careful consideration when interpreting our results. First, our identification of a CXCR3^+^ CD8^+^ NKT-like population does not resolve its full heterogeneity. CXCR3 marks diverse effector subsets—including Th1-biased CD4^+^ T cells, canonical CD8^+^ T_RM_, type I/II NKT cells, MAIT cells, γδT cells, and FOXP3^+^ regulatory T cells—each of which may respond differently to epithelial *Cxcl10* and exert distinct tissue-level effects.^37-40^ Second, the functional implications of sustaining these cytotoxic cells within the upper airway epithelium remain uncertain. Their transcriptional profile, defined by residency markers (*Itga1/*CD49a) and effector genes (*Gzma, Gzmb, Ccl5*), suggests they are primed memory-like cells capable of rapid antiviral defense. Indeed, emerging evidence positions CD49a as a compartmental discriminator for CD8^+^ T_RM_ cells in barrier tissues like skin, where it underpins rapid effector deployment (*Gzma, Gzmb, Ccl5, Nkg7, Cxcr3*) upon restimulation.^40^ By extension, our cells may afford protective immunosurveillance against reinfection, aligning with canonical T_RM_ benefits in respiratory models.^40,41^ However, their persistence may also potentiate maladaptive inflammation during recurrent insults. Distinguishing protective versus pathogenic roles will require studies incorporating lineage tracing, targeted depletion, cytokine blockade, and secondary-challenge paradigms.

These interpretive nuances extend to broader methodological constraints that shape our conclusions. For instance, we examined a single IAV strain and dose in young adult mice, leaving open questions about how aging, viral variants, or comorbidities (e.g., asthma or reflux) might reshape the MYC–CXCL10–CD8^+^ NKT pathway. Our conditional *Myc* knockout targeted KRT5^+^ basal cells broadly, without separating rare stem-like pools (e.g., KRT14^+^) from more differentiated ones, nor does it address potential compensation by other *Myc* paralogs over longer time scales. Moreover, although our analyses capture structural and molecular outcomes, they do not assess physiological consequences such as mucociliary clearance, airway biomechanics, or vagal reflex function.

Despite these limitations, our findings offer compelling clinical insights, revealing how a single viral hit can leave a lasting mark on airway tissue structure and local immune defenses. This mechanism is especially relevant to post-viral vagal neuropathy, chronic cough, voice loss (dysphonia), and related unexplained problems with laryngeal sensation and movement. In such conditions, an exaggerated or poorly resolved T_RM_-like response in the upper airway mucosa may sustain low-grade inflammation, skew epithelial differentiation, and perturb sensory circuits, thereby prolonging symptoms despite virologic recovery. Fundamentally, our results frame upper airway basal cells as key hubs linking repair to immune tuning, wherein MYC-driven proliferation and chemotactic cues (such as CXCL10-mediated immune influx) orchestrate balance between adaptive memory and maladaptive persistence.

To extend these findings, future studies should determine how repeated or mixed viral exposures recalibrate this epithelial–immune circuit and clarify when MYC-driven repair and CXCL10– CXCR3–mediated CD8^+^ NKT recruitment promote resilience versus precipitate maladaptive remodeling. Longitudinal approaches that integrate epithelial and immune profiling with direct assessments of neuroimmune function—such as vagal afferent activity, cough and laryngeal reflex sensitivity, and sensory nerve morphology—will be essential for establishing whether sustained cytotoxic surveillance disrupts sensory pathways in a manner that parallels human post-viral disease.

In summary, our study highlights basal *Myc* as a sentinel program that links epithelial repair to immune surveillance by coordinating epithelial growth, cell fate decisions, and chemokine-driven CD8^+^ NKT cell influx following viral injury. This coordinated program likely contributes to efficient viral control and the establishment of local memory but may also set the stage for persistent epithelial remodeling and neuroimmune dysregulation when dysbalanced.

## Methods

### Animals

All experimental procedures were performed in the American Association for Accreditation of Laboratory Animal Care (AAALAC)-certified laboratory animal facility at the University of California, San Diego (UCSD). All animal husbandry and experiments were conducted under approved Institutional Animal Care and Use Committee (IACUC) guidelines. Wild-type C57BL/6 (JAX 000664), *Krt5*^*CreERT2*^ (JAX 029155), *Ascl1*^*CreERT2*^ (JAX 012882), and *Rosa*^*lxl-tdTomato*^ (*Ai14*, JAX 007914) lines were purchased from the Jackson lab. *Myc-flox* (MMRRC 036778) line was gifted from Rob Signer at UCSD. All the *cre* lines we used in this study were kept in C57BL/6 background. All *cre* driver lines in heterozygous form are viable and fertile, and no abnormal phenotypes were detected. Both male and female mice were used in the experiment. Adult mice were 8-12 weeks of age for all experiments. The experiments were not randomized, and the investigators were not blinded to allocation during experiments and outcome assessment.

### Influenza A virus (IAV) infection

Influenza strain A/H1N1/PR/8 strain was obtained from ATCC (VR-95PQ). Briefly, mice were anesthetized with 3.5% isoflurane for 15 min until bradypnea was observed. Virus dissolved in 35 µl of PBS was pipetted onto the nostrils of anesthetized mice, whereupon they aspirated the fluid directly into their airways. Control and experimental groups were infected simultaneously in the same cohort by the same investigator so that direct comparison between groups was justified and appropriate.

### Tissue collection and immunofluorescence

Mice were euthanized by CO2 inhalation followed by transcardial perfusion with PBS to remove circulating blood. The larynx-trachea was isolated and fixed overnight in 1%PFA or immediately embedded in OCT, flash frozen using 2-Methylbutane and liquid nitrogen and stored at –80°C. Using a cryostat, the larynx-trachea was then sectioned (12 µm) and stored at –20°C. Unfixed/lightly fixed tissue was then washed in PBS for 5mins to remove OCT, heated to boiling in 10 mM citrate buffer (pH 9) for antigen retrieval and treated with 0.5% Triton X-100 in PBS for 15mins. All sections were processed for immunostaining following a standard protocol.^42^ *Ascl1* and *Krt5* recombination was induced via Tamoxifen IP administration with dose of 100mg/kg for either two or three consecutive days. All primary and secondary antibodies used are listed in TableS2. Primary antibodies were applied overnight at 4°C, while secondary antibodies were applied for 1hr at room temperature. Sections were incubated with DAPI (1:1000 ratio) for 10 min at RT. Slides were mounted and coverslipped with Prolong Diamond mounting media (Fisher P36970), cured flat at room temperature in the dark for 24 h, and stored at 4°C. Each experiment was replicated at least twice for all timepoints and targets assessed.

### Tissue processing and cell sorting

Whole upper airway (larynx-trachea) was mechanically dissociated by mincing tissue with razor blades in solution containing 5 ml of RPMI 1640 (Thermo Scientific) with 10% fetal bovine serum, 1 mM HEPES (Life Technology), 1 mM MgCl_2_ (Life Technology), 1 mM CaCl_2_ (Sigma), 0.5 mg ml^−1^ collagenase D and type I/Dispase (Roche), and 0.25 mg DNase I (Roche). Minced tissue was then digested by shaking at around 150 rpm for 30 min at 37 °C. Following incubation, upper airway pieces were mechanically dissociated further by straining through a MACS 70 µm filter. Red blood cells were removed by the addition of 1 ml of RBC lysis buffer (BioLegend) to each tube and incubation at room temperature for 1 min. Single-cell suspensions were pelleted (1,500 rpm, 4 °C, 5 min) and stained with Fc blocking antibody (5 mg ml^−1^, BD). For FACS, the following antibodies were used: (Immune) 1:1000 BV510-conjugated anti-CD45 (BioLegend, no. 103138), (Epithelial) 1:1000 APC-FITC-conjugated anti-Epcam (BioLegend, no. 118214), and (Endothelial) 1:1000 PE-conjugated anti-CD31 (BioLegend, no. 102508). Cells were then stained using live/dead dye (CD11b: Brilliant Violet 450, BioLegend, no. 75-0112-U025) before being resuspended in 2% FBS + 1:2000 DAPI. All FACS sorting for scRNA-seq library preparation was done on a BD FACSAria Fusion Sorter (BD Biosciences) analyzer with three lasers (405, 488 and 640 nm) at the Flow Cytometry Core at VA San Diego Health Care System.

### Bulk RNA-seq and data analysis

Total RNA from 10wk old adult WT upper airway (larynx/trachea) tissue was extracted as described above. cDNA libraries were constructed using Illumina TruSeq RNA Library Prep Kit V2 (Illumina) and sequenced on the HiSeq4000 platform (Illumina) at the Institute for Genomic Medicine (IGM) at UCSD. FASTQ files were aligned to the mouse reference genome (mm10) by using Bowtie2^43^ with default settings. Differential gene expression analysis was performed using Cufflinks.^44^ P-values were adjusted by the Benjamini-Hochberg method to control False Discovery Rate (FDR) at 0.05. Heatmaps and volcano plots were generated by ggplot2 (version 3.3.2) using RStudio. (v. 2024.04.0 Build 735) running R 4.3.3 (R Core Team). Once exclusively differentially expressed genes were identified, we performed tests of enrichment using Gene Ontology (GO) annotations utilizing Enrichr (v2024)^45^ as previously described.^46^

### 10x Chromium scRNAseq and data analysis

Single-cell RNA sequencing (scRNA-seq) was performed using the 10× Genomics Chromium platform as previously described.^10^ Briefly, dissociated cell suspensions were loaded onto the Chromium Controller to generate single-cell gel bead emulsions, followed by cDNA synthesis and library preparation using the Chromium Single Cell 3′ v3 kit (10X Genomics, Pleasanton, CA). Libraries were sequenced on an Illumina NovaSeq 6000, and raw reads were processed with Cell Ranger (v3.0.2, 10× Genomics) using the mouse reference genome (GRCm38) to generate single-cell gene barcode matrices. Influenza A Virus complete gene sequence was imputed into the reference genome for *in silico* transcript identification (TableS1). Downstream analyses—including quality control, normalization, clustering, and differential expression—were performed in Seurat (v4.0.5)^47,48^ following the previously mentioned computational workflow.^10^ Unless otherwise noted, identical parameters and filtering thresholds were applied. To account for technical and biological covariates, cell cycle phase scores (S and G2/M), mitochondrial transcript percentage, and gene feature counts (nFeature_RNA) were incorporated into the dataset and regressed during data scaling to minimize their confounding effects on clustering.

To infer and quantify cell–cell signaling, we applied the CellChat R package (v1.x) to the processed single-cell transcriptomic data. Ligand–receptor interactions and their associated cofactors were evaluated using the curated CellChatDB database (mouse version, ∼2,021 validated interactions).^27^ Communication networks were represented as directed, weighted graphs, and visualized using hierarchical plots, circle plots, bubble plots, and chord diagrams. Centrality metrics (e.g. in-degree, out-degree, betweenness) were computed to determine dominant signaling sources, targets, and mediators. Pattern recognition and manifold learning approaches were applied to identify global communication patterns and context-specific signaling differences across conditions.

To assess pathway-level activity across our IAV-enriched epithelial subpopulations, we applied the irGSEA R package (v2.1.5) for rank-based, single-cell gene set enrichment analysis.^49^ Gene sets were derived from MSigDB (Hallmark and KEGG collections) (version 7.4.1). Normalized expression matrices from Seurat objects were imported into irGSEA, and enrichment scores were calculated using the AUCell (version 1.14.0), UCell (version 1.1.0), singscore (version 1.12.0), GSVA (version 1.40.1) and Viper (version 1.32.0) algorithms implemented within the package. Scores were computed on a per-cell basis and summarized by cell type and condition. Differential pathway enrichment was visualized as violin and dot plots to highlight compartment-specific activation patterns. All parameters followed default settings in irGSEA unless otherwise specified.

Cellular lineage and differentiation trajectories were reconstructed using Monocle3 (1.4.26) to infer dynamic transcriptional transitions within epithelial IAV-enriched subpopulations.^50^ Briefly, Seurat-processed data were converted to a *cell_data_set* object using the *as*.*cell_data_set()* function, retaining normalized expression values and annotated metadata. Dimensionality reduction was performed using UMAP and PCA embeddings previously generated in Seurat. Cells were ordered in pseudotime using *learn_graph()* and *order_cells()*, with root nodes defined by basal populations. Pseudotime-associated genes were identified using *graph_test()*, and trajectory-dependent gene expression patterns were visualized using *plot_cells()* functions. Default Monocle3 parameters were used unless otherwise noted.

### RNAscope in situ hybridization

All staining procedures were performed using the RNAscope Fluorescent Multiplex Kit V2 (Advanced Cell Diagnostics, no. 323100) following the manufacturer’s instructions. The following probes from Advanced Cell Diagnostics were used: Mm-*IAV* (no. 521181), Mm-*Myc* (no. 413451), Mm-*Cxcl10* (no. 408921), Mm-*tdTomato* (no. 317041), Mm-*Gzmb* (no. 490191), Mm-*Ccl5* (no. 469601).

### Cell proliferation and apoptosis

Cell proliferation was assessed using the Click-iT™ EdU Cell Proliferation Kit (C10337, Invitrogen), which detects DNA synthesis through copper-catalyzed azide–alkyne cycloaddition between incorporated 5-ethynyl-2′-deoxyuridine (EdU) and a fluorescent azide probe. Adult mice received 1 mL of a 400 µM EdU solution (Invitrogen, diluted in PBS) by intraperitoneal injection, and upper airway tissues were collected 1 hour later. Samples were fixed in 1% paraformaldehyde (PFA) overnight at 4°C, washed in PBS for 30 minutes, and embedded in OCT before cryosectioning and EdU detection according to the manufacturer’s protocol.

Apoptosis was measured using the Click-iT™ Plus TUNEL Assay (C10245, Invitrogen), which labels DNA strand breaks via terminal deoxynucleotidyl transferase (TdT)–mediated incorporation of alkyne-modified nucleotides, followed by fluorescent azide detection using the same click-chemistry–based reaction. Tissue preparation, fixation, and embedding were identical to those described for EdU staining, and TUNEL labeling was performed following the manufacturer’s instructions.

### Lineage tracing

Tamoxifen dissolved in corn oil was administered via intraperitoneal injection at 25 mg/kg once daily for three consecutive days.

### Cell count quantification

All cell counts and area measurements were quantified at 20× magnification from 3–4 whole-mount coronal images spanning the pharyngolaryngeal-to-tracheobronchial axis (Fig. S1B). Images were analyzed using an automated pipeline in QuPath (v0.6.0). Briefly, the epithelium was manually annotated as a region of interest (ROI) and subdivided into regional compartments, including tracheal surface epithelium (SE), subglottic SE, supraglottic SE, and submucosal glands (SMG). For marker-positive cell counts or total feature area (µm2), the QuPath *create_threshold* function was applied within each annotated epithelial ROI to accurately label the target signal. The *cell_detection* function was then used within the same ROI to identify DAPI^+^ nuclei and determine total cell numbers. Threshold-defined feature counts were normalized to DAPI^+^ counts to calculate the percentage of positive cells; alternatively, threshold-defined feature area was recorded within each ROI and reported as total area (µm2) or expressed in arbitrary units (AU) as indicated.

### Statistics

All analyses were performed using GraphPad Prism 10. Data were first assessed for normality (Shapiro–Wilk) and homogeneity of variance (F test) to determine whether parametric or non-parametric tests were appropriate. For normally distributed data with equal variance, statistical comparisons used t-tests (two groups) or one-way/two-way ANOVA (multiple groups or multiple variables), followed by the appropriate post-hoc corrections (Dunnett’s, Tukey’s, or Fisher’s LSD) as indicated in figure legends. When normality or variance assumptions were not met, non-parametric equivalents were applied. Results are reported as mean ± SE, and significance was defined as p ≤ 0.05.

## Data and code availability

Bulk and single cell RNA-seq data have been deposited at the Gene Expression Omnibus (GEO) and are publicly available as of the date of publication. All data generated supporting the findings of this study are available in the manuscript. Further information is available from the lead author upon reasonable request. This paper does not report original code.

## Acknowledgements

The authors thank Sun lab members for discussions. UCSD Microscopy Core was supported by NINDS-P30NS047101. This work was supported by grants NIH NHLBI R01 AT011676-01 (to X.S.), 1R01 HL160019-01 (to X.S.), NIH NIDCD F32 DC021634-01 (to A.G.F.) and ASLHF (to A.G.F.).

## Author contributions

Conceptualization: A.G.F, X.S. Methodology: A.G.F. Investigation: A.G.F., L.X., B.P., N.K., J.V. Visualization: A.G.F. Funding acquisition: A.G.F., X.S. Supervision: X.S. Writing – original draft: A.G.F. Writing – review & editing: A.G.F., X.S.

## Competing interests

The authors declare no competing interests.

## SUPPLEMENTAL FIGURES

**Fig. S2.**
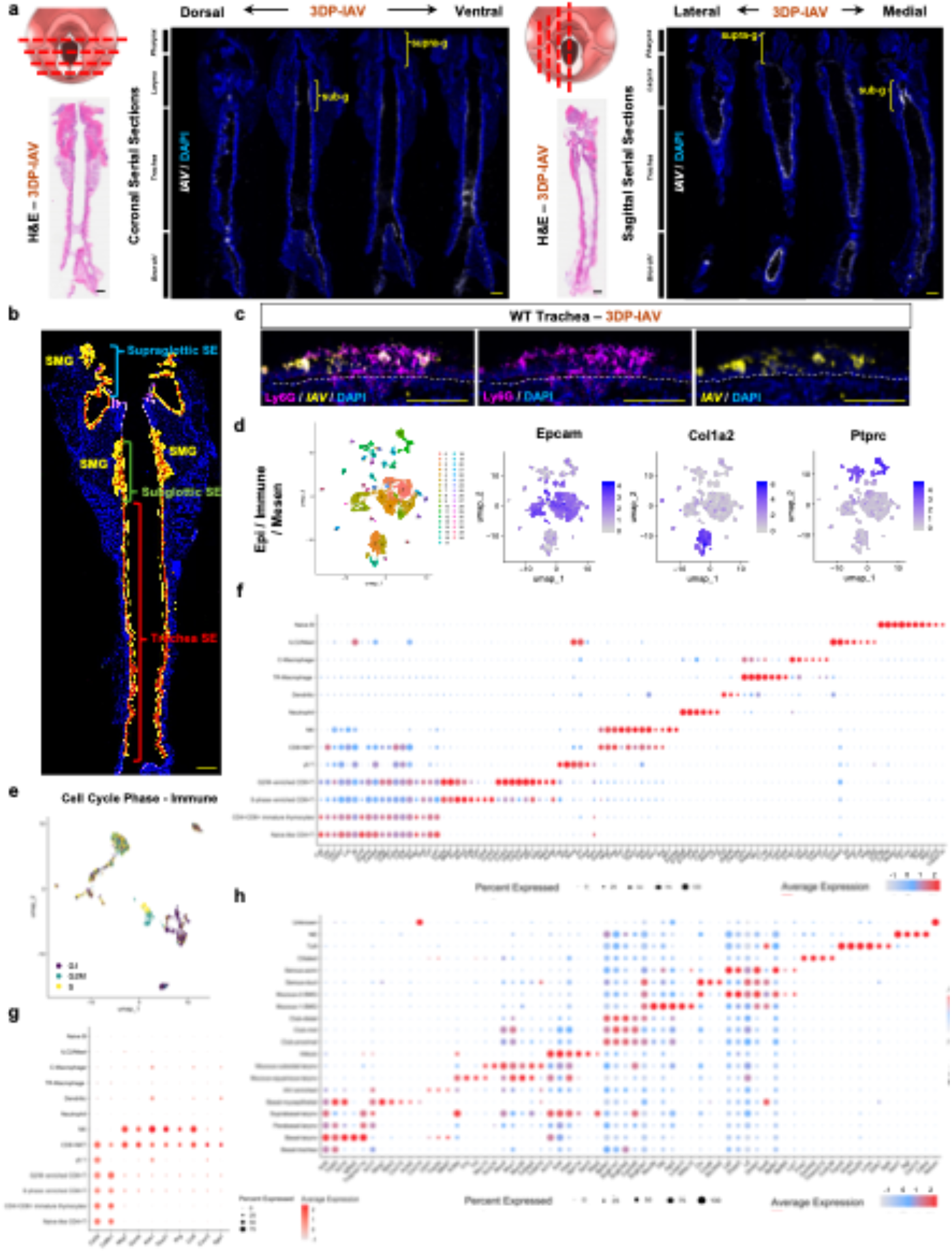
Validation of viral transcript integration and MYC expression in IAV-enriched airway epithelium. **a**, Data-integrity check: UMAPs before and after removing the synthetic IAV transcript show identical clustering and annotations, indicating no distortion from viral transcript integration. **b**, Dot plot showing *Myc* and *Mycl* expression across epithelial subtypes at 7DP-IAV; *Mycn* not found. Circle size reflects the percentage of cells expressing each gene; color indicates average expression level. **c**, Quantification of the proportion of *Myc*^+^ cells within IAV-enriched epithelial populations. **d**, H&E-stained coronal sections of tracheal and subglottic epithelium from control and Krt5^creER^;Myc conditional mutants at 21DP-SALINE. **e**, Body weight measurements of male and female control versus Krt5^creER^;Myc mice at baseline (Day 0) and at 7DP-IAV; one-way ANOVA, Tukey’s post-hoc analysis to compare group means, mean ± SEM. **f**, RNA *in situ* hybridization of IAV transcripts (magenta) in tracheal sections from control and Krt5^creER^;Myc mice at 7DP-IAV and 10DP-IAV, showing comparable viral clearance kinetics between genotypes. DP days post.

**Fig. S2.**
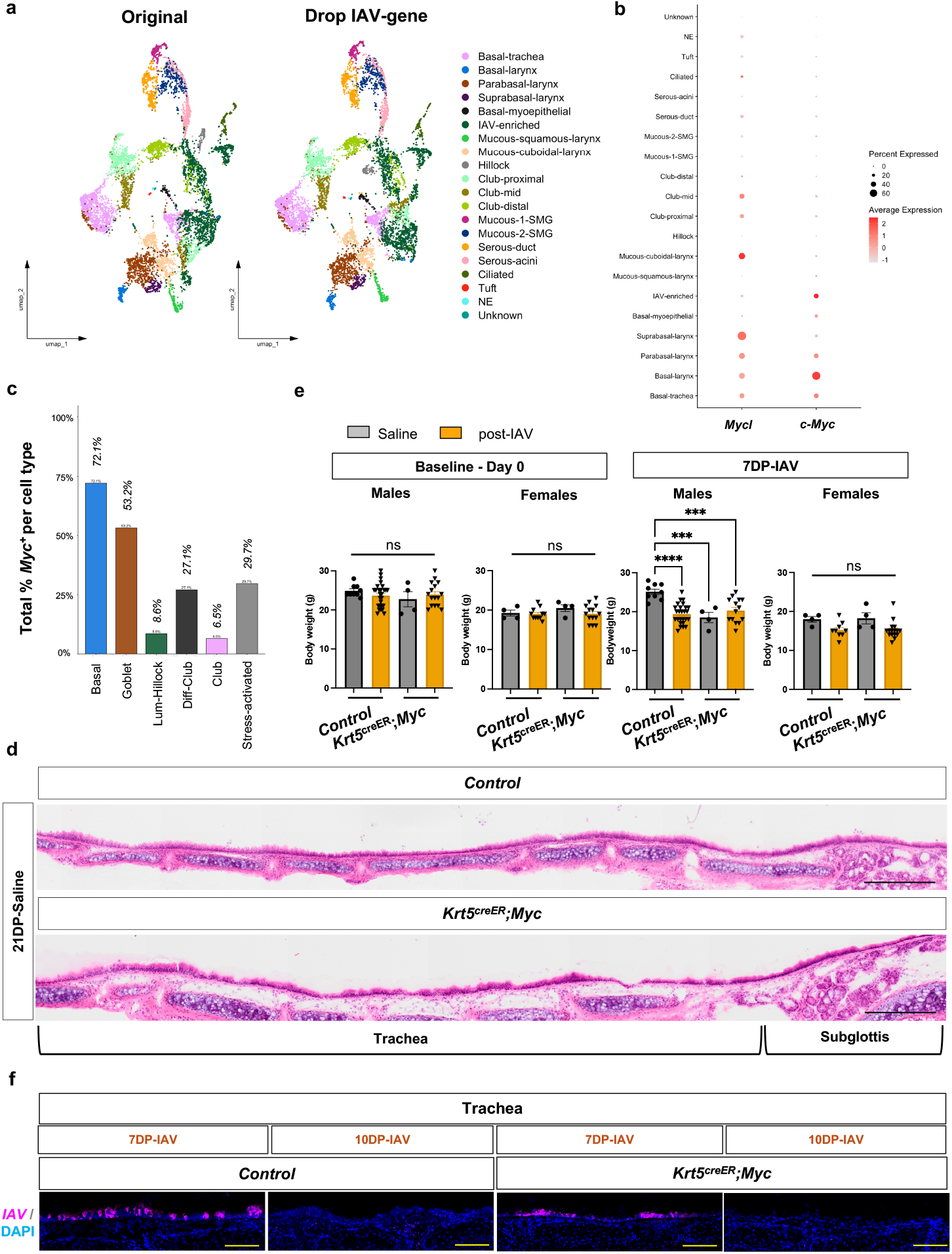
Validation of viral transcript integration and MYC expression in IAV-enriched airway epithelium. **a**, Data-integrity check: UMAPs before and after removing the synthetic IAV transcript show identical clustering and annotations, indicating no distortion from viral transcript integration. **b**, Dot plot showing *Myc* and *Mycl* expression across epithelial subtypes at 7DP-IAV; *Mycn* not found. Circle size reflects the percentage of cells expressing each gene; color indicates average expression level. **c**, Quantification of the proportion of *Myc*^+^ cells within IAV-enriched epithelial populations. **d**, H&E-stained coronal sections of tracheal and subglottic epithelium from control and Krt5^creER^;Myc conditional mutants at 21DP-SALINE. **e**, Body weight measurements of male and female control versus Krt5^creER^;Myc mice at baseline (Day 0) and at 7DP-IAV; one-way ANOVA, Tukey’s post-hoc analysis to compare group means, mean ± SEM. **f**, RNA *in situ* hybridization of IAV transcripts (magenta) in tracheal sections from control and Krt5^creER^;Myc mice at 7DP-IAV and 10DP-IAV, showing comparable viral clearance kinetics between genotypes. DP days post..

**Fig. S3.**
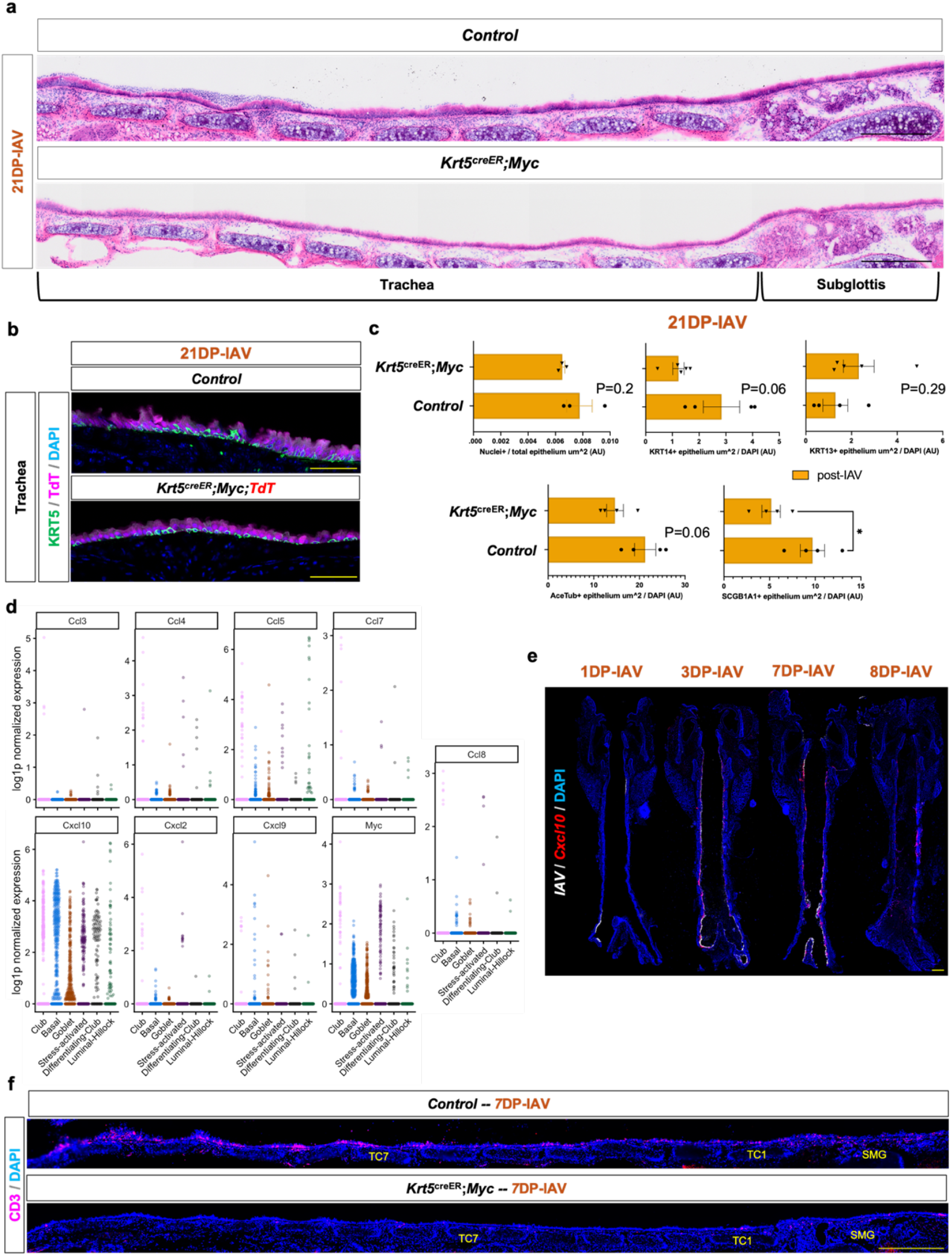
MYC regulates epithelial immune signaling and CD3^+^ cell accumulation following IAV infection. **a**, H&E-stained coronal sections of tracheal and subglottic epithelium from control and Krt5^creER^;Myc conditional mutants at 21DP-IAV. **b**, Immunofluorescence imaging of KRT5 (green) and TdTomato lineage label (magenta) at 21DP-IAV. **c**, Quantification of total nuclei and lineage-defined epithelial compartments at 21DP-IAV, normalized to total epithelial area (um^2) and (DAPI^+^) counts, respectively. **d**, Violin plots of immune-related transcripts from scRNA-seq across IAV-enriched epithelial clusters, showing differential expression of *Cd3d, Ccl4, Ccl5, Ccl7, Cd8b, Cxcl10, Cxcl2, Cxcl9*, and *Myc* at 7DP-IAV. **e**, Dynamic *Cxcl10* expression tracks sites of active IAV replication. **f**, Immunofluorescence at 7DP-IAV showing reduced CD3^+^ cells to upper respiratory airway in *Myc* mutant mice compared to controls. Scale bars, 500 µm. *P < 0.05, **P < 0.005, ***P<0.0005; ns, not significant. DP days post.

## SUPPLEMENTAL TABLES

### Influenza A Virus (A/Puerto Rico/8/1934(H1N1)) reference gene sequence

ATGAAGGCAAACCTACTGGTCCTGTTATGTGCACTTGCAGCTGCAGATGCAGACACAATATGTATAGGCTACCATGCGAACAATTCAACCGACACTGTTGACACAGTGCTCGAGAAGAATGTGACAGTGACACACTCTGTTAACCTGCTCGAAGACAGCCACAACGGAAAACTATGTAGATTAAAAGGAATAGCCCCACTACAATTGGGGAAATGTAACATCGCCGGATGGCTCTTGGGAAACCCAGAATGCGACCCACTGCTTCCAGTGAGATCATGGTCCTACATTGTAGAAACACCAAACTCTGAGAATGGAATATGTTATCCAGGAGATTTCATCGACTATGAGGAGCTGAGGGAGCAATTGAGCTCAGTGTCATCATTCGAAAGATTCGAAATATTTCCCAAAGAAAGCTCATGGCCCAACCACAACACAACCAAAGGAGTAACGGCAGCATGCTCCCATGCGGGGAAAAGCAGTTTTTACAGAAATTTGCTATGGCTGACGGAGAAGGAGGGCTCATACCCAAAGCTGAAAAATTCTTATGTGAACAAGAAAGGGAAAGAAGTCCTTGTACTGTGGGGTATTCATCACCCGTCTAACAGTAAGGATCAACAGAATATCTATCAGAATGAAAATGCTTATGTCTCTGTAGTGACTTCAAATTATAACAGGAGATTTACCCCGGAAATAGCAGAAAGACCCAAAGTAAGAGATCAAGCTGGGAGGATGAACTATTACTGGACCTTGCTAAAACCCGGAGACACAATAATATTTGAGGCAAATGGAAATCTAATAGCACCAAGGTATGCTTTCGCACTGAGTAGAGGCTTTGGGTCCGGCATCATCACCTCAAACGCATCAATGCATGAGTGTAACACGAAGTGTCAAACACCCCTGGGAGCTATAAACAGCAGTCTCCCTTTCCAGAATATACACCCAGTCACAATAGGAGAGTGCCCAAAATACGTCAGGAGTGCCAAATTGAGGATGGTTACAGGACTAAGGAACATTCCGTCCATTCAATCCAGAGGTCTATTTGGAGCCATTGCCGGTTTTATTGAAGGGGGATGGACTGGAATGATAGATGGATGGTACGGTTATCATCATCAGAATGAAC AGGGATCAGGCTATGCAGCGGATCAAAAAAGCACACAAAATGCCATTAACGGGATTACAAACAAGGTGAACTCTGTTATCGAGAAAATGAACATTCAATTCACAGCTGTGGGTAAAGAATTCAACAAATTAGAAAAAAGGATGGAAAATTTAAATAAAAAAGTTGATGATGGATTTCTGGACATTTGGACATATAATGCAGAATTGTTAGTTCTACTGGAAAATGAAAGGACTCTGGATTTCCATGACTCAAATGTGAAGAATCTGTATGAGAAAGTAAAAAGCCAATTAAAGAATAATGCCAAAGAAATCGGAAATGGATGTTTTGAGTTCTACCACAAGTGTGACAATGAATGCATGGAAAGTGTAAGAAATGGGACTTATGATTATCCCAAATATTCAGAAGAGTCAAAGTTGAACAGGGAAAAGGTAGATGGAGTGAAATTGGAATCAATGGGGATCTATCAGATTCTGGCGATCTACTCAACTGTCGCCAGTTCACTGGTGCTTTTGGTCTCCCTGGGGGCAATCAGTTTCTGGATGTGTTCTAATGGATCTTTGCAGTGCAGAATATGCATCTGA

### Clustering fidelity analysis results before and after exclusion of synthetic IAV-gene ARI:NMI Confusion Matrix

ARI:NMI Confusion Matrix
|  |  | DropIAV |  |  |  |  |
| --- | --- | --- | --- | --- | --- | --- |
| Original | 1 | 10 | 11 | 12 | 13 | 14 |
| 1 | 1123 | 0 | 0 | 0 | 0 | 0 |
| 10 | 0 | 342 | 0 | 0 | 0 | 0 |
| 11 | 0 | 0 | 269 | 0 | 0 | 0 |
| 12 | 0 | 0 | 0 | 162 | 0 | 0 |
| 13 | 0 | 0 | 0 | 0 | 148 | 0 |
| 14 | 0 | 0 | 0 | 0 | 0 | 131 |

### Percentage of cells expressing “IAV reference gene” within each major cell compartment and cell type

**Percentage of cells expressing "IAV reference gene" within each major cell compartment and cell type**
| Gene | Pharyngolaryngeal % | Tracheobronchial % | Submucosal % | IAV-enriched % |
| --- | --- | --- | --- | --- |
| *Tmprss11a+* | 64.87576 | 10.6769 | 2.629544 | 12.24242 |
| *Nkx2-1+* | 15.17797 | 60.36288 | 33.79737 | 33.57576 |
| *Dmbt1+* | 35.93015 | 42.63782 | 90.719258 | 26.9697 |

| Gene | CellType | Total cells_Control | Total cells_post-IAV | CellType expressing IAV-gene % |
| --- | --- | --- | --- | --- |
| IAV-gene | Club-proximal | 486 | 170 | 10.21 |
| IAV-gene | Club-distal | 290 | 121 | 6.81 |
| IAV-gene | Club-mid | 291 | 120 | 10.95 |
| IAV-gene | Basal-trachea | 847 | 282 | 4.96 |
| IAV-gene | Serous-acini | 328 | 73 | 3.49 |
| IAV-gene | Mucous-1-SMG | 91 | 71 | 11.73 |
| IAV-gene | Ciliated | 67 | 64 | 28.24 |
| IAV-gene | Mucous-cuboidal-larynx | 208 | 241 | 16.48 |
| IAV-gene | Mucous-2-SMG | 191 | 117 | 7.14 |
| IAV-gene | Parabasal-larynx | 247 | 387 | 13.56 |
| IAV-gene | Serous-duct | 152 | 194 | 14.45 |
| IAV-gene | Mucous-squamous-larynx | 35 | 115 | 38.00 |
| IAV-gene | IAV-enriched | 152 | 1498 | 32.12 |
| IAV-gene | Suprabasal-larynx | 53 | 71 | 8.87 |
| IAV-gene | Basal-larynx | 73 | 59 | 5.30 |
| IAV-gene | Hillock | 56 | 40 | 7.29 |
| IAV-gene | Basal-myoepithelial | 34 | 42 | 15.79 |
| IAV-gene | Tuft | 20 | 0 | 0.00 |
| IAV-gene | NE | 5 | 0 | 0.00 |
| IAV-gene | Unknown | 1 | 6 | 14.29 |

### Percentage of cells in S or G2M phase for each epithelial cluster

| CellType | total_cells | cycling_cells | percentage_cycling |
| --- | --- | --- | --- |
| Basal-trachea | 1129 | 300 | 26.572188 |
| Basal-larynx | 132 | 84 | 63.636364 |
| Parabasal-larynx | 634 | 129 | 20.347003 |
| Suprabasal-larynx | 124 | 9 | 7.258065 |
| Basal-myoepithelial | 76 | 18 | 23.684211 |
| IAV-enriched | 1650 | 1197 | 72.545455 |
| Mucous-squamous-larynx | 150 | 27 | 18 |
| Mucous-cuboidal-larynx | 449 | 48 | 10.690423 |
| Hillock | 96 | 36 | 37.5 |
| Club-proximal | 656 | 133 | 20.27439 |
| Club-mid | 411 | 56 | 13.625304 |
| Club-distal | 411 | 218 | 53.041363 |
| Mucous-1-SMG | 162 | 15 | 9.259259 |
| Mucous-2-SMG | 308 | 37 | 12.012987 |
| Serous-duct | 346 | 47 | 13.583815 |
| Serous-acini | 401 | 99 | 24.688279 |
| Ciliated | 131 | 55 | 41.984733 |
| Tuft | 20 | 4 | 20 |
| NE | 5 | 0 | 0 |
| Unknown | 7 | 0 | 0 |

### REAGENT or RESOURCE

| REAGENT or RESOURCE | SOURCE | IDENTIFIER |
| --- | --- | --- |
| **Antibodies** |  |  |
| Rabbit-KRT5 | Abcam | Cat#ab52635 |
| Rabbit, anti-KRT13 | Abcam | Cat#ab92551 |
| Mouse, anti-KRT14 | Millipore | Cat#CBL197 |
| Rabbit, anti-KRT17 | Santa Cruz Biotechnology | Cat#sc-393002 |
| Mouse-aSMA-FITC | Millipore Sigma | Cat#F3777 |
| Rat-CD3 | Abcam | Cat#ab11089 |
| Rat-Ly6G | BioLegend | Cat#127601 |
| Mouse-RFP | Abcam | Cat#ab125244 |
| Rabbit-DsRed | Takara Clontech | Cat#632496 |
| Mouse-FOXJ1 | eBioscience/Fisher | Cat#5013931 |
| Mouse-Acetylated tubulin | Sigma | Cat#T7451 |
| Rabbit-SCGB1A1 | Abcam | Cat#ab213203 |
| Goat anti-Rabbit IgG Alexa Flour 488 | ThermoFisher | Cat#A27034 |
| Goat anti-mouse IgG, Cy3<sup>TM</sup> | Jackson Immuno Research | Cat#115-166-003 |
| Goat anti-rat IgG, Cy3<sup>TM</sup> | Jackson Immuno Research | Cat#112-165-003 |
| BV510-conjugated anti-CD45 | BioLegend | Cat#103138 |
| APC-FITC-conjugated anti-Epcam | BioLegend | Cat#118214 |
| PE-conjugated anti-CD31 | BioLegend | Cat#102508 |
| CD11b: Brilliant Violet 450 | BioLegend | Cat#75-0112-U025 |
| Mm-IAV | Advanced Cell Diagnostics | Cat#521181 |
| Mm-Myc | Advanced Cell Diagnostics | Cat#413451 |
| Mm-Cxcl10 | Advanced Cell Diagnostics | Cat#408921 |
| Mm-tdTomato | Advanced Cell Diagnostics | Cat#317041 |
| Mm-Gzmb | Advanced Cell Diagnostics | Cat#490191 |
| Mm-Ccl5 | Advanced Cell Diagnostics | Cat#469601 |
| **Bacterial and virus strains** |  |  |
| Influenza strain A/H1N1/PR/8 | ATCC | VR-95PQ |
| **Chemicals, peptides, and recombinant proteins** |  |  |
| Tamoxifen | Sigma-Aldrich | Cat#T-5648-1G |
| RPMI 1640 Medium | Thermo Scientific | Cat#11875093 |
| 1 mM HEPES | Life Technology | Cat#15630080 |
| Dispase | Roche | Cat#4942086001 |
| DNase I | Roche | Cat#11284932001 |
| 1 mM CaCl<sub>2</sub> | Sigma-Aldrich | Cat#21115 |
| 1 mM MgCl<sub>2</sub> | Sigma-Aldrich | Cat#20303M |
| Collagenase D | Roche | Cat#11088858001 |
| RBC Lysis Buffer | BioLegend | Cat#420301 |
| **Critical commercial assays** |  |  |
| RNAscope® Multiplex Fluorescent Reagent Kit V2 | Advanced Cell Diagnostics | Cat#323100 |
| Click-it TUNEL Alexa Flour 488 | ThermoFisher | Cat#C10245 |
| Click-it EdU Alexa Flour 488 | ThermoFisher | Cat#C10337 |
| **Deposited data** |  |  |
| Single cell RNA-seq data for mouse larynx/trachea | Gene Expression Omnibus | n/a |
| bulk RNA-seq data for mouse larynx/trachea | Gene Expression Omnibus | n/a |
| **Experimental models: Organisms/strains** |  |  |
| Mouse: Wildtype C57BL/6J | Jackson laboratory | JAX: 000664 |
| Mouse: Rosa-*lxl*-tdTomato (Ai14) | Jackson laboratory | JAX: 007914 |
| Mouse: Ascl1-creERT2 | Jackson laboratory | JAX: 012882 |
| Mouse: Krt5-creERT2 | Jackson laboratory | JAX: 029155 |
| Mouse: Myc-flox | Gifted from Rob Signer at UCSD | MMRRC 036778 |
| **Software and algorithms** |  |  |
| RStudio (v2024.12.1+563) | RStudio Team | http://www.rstudio.com/ |
| R (v4.5.0) | R Core Team | https://www.R-project.org/ |
| Cell Ranger (v3.0.2) | 10x Genomics | https://support.10xgenomics.com/single-cell-gene-expression/software/pipelines/latest/what-is-cell-ranger |

## Notes

### Competing Interest Statement

The authors have declared no competing interest.

### Summary of Updates

Updated text edits, and figure panel rearrangement.

## References

1 An, R. et al. Single-Cell View into the Role of Microbiota Shaping Host Immunity in the Larynx. iScience (2024).

2 Jette, M. E., Dill-McFarland, K. A., Hanshew, A. S., Suen, G. & Thibeault, S. L. The human laryngeal microbiome: effects of cigarette smoke and reflux. Sci Rep 6, 35882 (2016). 10.1038/srep35882

3 Thibeault, S. L., Rees, L., Pazmany, L. & Birchall, M. A. At the crossroads: mucosal immunology of the larynx. Mucosal Immunol 2, 122–128 (2009). 10.1038/mi.2008.82

4 Niec, R. E., Rudensky, A. Y. & Fuchs, E. Inflammatory adaptation in barrier tissues. Cell 184, 3361–3375 (2021). 10.1016/j.cell.2021.05.036

5 Hewitt, R. J. & Lloyd, C. M. Regulation of immune responses by the airway epithelial cell landscape. Nat Rev Immunol 21, 347–362 (2021). 10.1038/s41577-020-00477-9

6 Yang, J. & Yan, H. Mucosal epithelial cells: the initial sentinels and responders controlling and regulating immune responses to viral infections. Cell Mol Immunol 18, 1628–1630 (2021). 10.1038/s41423-021-00650-7

7 Chen, X. et al. Host Immune Response to Influenza A Virus Infection. Front Immunol 9, 320 (2018). 10.3389/fimmu.2018.00320

8 King, R. E. et al. A Novel In Vivo Model of Laryngeal Papillomavirus-Associated Disease Using Mus musculus Papillomavirus. Viruses 14 (2022). 10.3390/v14051000

9 King, R. E. et al. The Larynx is Protected from Secondary and Vertical Papillomavirus Infection in Immunocompetent Mice. The Laryngoscope (2023).

10 Foote, A. G. & Sun, X. Cellular Heterogeneity and Patterning Strategies as Revealed by Upper Respiratory Epithelium Single Cell Atlas. iScience, 112845 (2025).

11 Lin, B. et al. Airway hillocks are injury-resistant reservoirs of unique plastic stem cells. Nature (2024). 10.1038/s41586-024-07377-1

12 Jin, X. et al. Global burden of upper respiratory infections in 204 countries and territories, from 1990 to 2019. EClinicalMedicine 37, 100986 (2021). 10.1016/j.eclinm.2021.100986

13 Lee, B. & Woo, P. Chronic cough as a sign of laryngeal sensory neuropathy: diagnosis and treatment. Annals of Otology, Rhinology & Laryngology 114, 253–257 (2005).

14 Rees, C. J., Henderson, A. H. & Belafsky, P. C. Postviral vagal neuropathy. Ann Otol Rhinol Laryngol 118, 247–252 (2009). 10.1177/000348940911800402

15 Garcia-Vicente, P. et al. Chronic cough in post-COVID syndrome: Laryngeal electromyography findings in vagus nerve neuropathy. PLoS One 18, e0283758 (2023). 10.1371/journal.pone.0283758

16 Amin, M. R. & Koufman, J. A. Vagal neuropathy after upper respiratory infection: a viral etiology? Am J Otolaryngol 22, 251–256 (2001). 10.1053/ajot.2001.24823

17 Jeyakumar, A., Brickman, T. M. & Haben, M. Effectiveness of amitriptyline versus cough suppressants in the treatment of chronic cough resulting from postviral vagal neuropathy. Laryngoscope 116, 2108–2112 (2006). 10.1097/01.mlg.0000244377.60334.e3

18 Bhatt, N. K., Pipkorn, P. & Paniello, R. C. Association between upper respiratory infection and idiopathic unilateral vocal fold paralysis. Annals of Otology, Rhinology & Laryngology 127, 667–671 (2018).

19 Smith, A. M. & Perelson, A. S. Influenza A virus infection kinetics: quantitative data and models. Wiley Interdiscip Rev Syst Biol Med 3, 429–445 (2011). 10.1002/wsbm.129

20 Bin, N. R. et al. An airway-to-brain sensory pathway mediates influenza-induced sickness. Nature (2023). 10.1038/s41586-023-05796-0

21 Wang, W. et al. Stress keratin 17 enhances papillomavirus infection-induced disease by downregulating T cell recruitment. PLoS Pathog 16, e1008206 (2020). 10.1371/journal.ppat.1008206

22 Rudd, J. M. et al. Neutrophils Induce a Novel Chemokine Receptors Repertoire During Influenza Pneumonia. Front Cell Infect Microbiol 9, 108 (2019). 10.3389/fcimb.2019.00108

23 Krenz, B., Lee, J., Kannan, T. & Eilers, M. Immune evasion: An imperative and consequence of MYC deregulation. Mol Oncol 18, 2338–2355 (2024). 10.1002/1878-0261.13695

24 Pelengaris, S. & Khan, M. The many faces of c-MYC. Arch Biochem Biophys 416, 129–136 (2003). 10.1016/s0003-9861(03)00294-7

25 Dhanasekaran, R. et al. The MYC oncogene— the grand orchestrator of cancer growth and immune evasion. Nature reviews Clinical oncology 19, 23–36 (2022).

26 Portal, C., Wang, Z., Scott, D. K., Wolosin, J. M. & Iomini, C. The c-Myc oncogene maintains corneal epithelial architecture at homeostasis, modulates p63 expression, and enhances proliferation during tissue repair. Invest Ophth Vis Sci 63, 3–3 (2022).

27 Jin, S. et al. Inference and analysis of cell-cell communication using CellChat. Nat Commun 12, 1088 (2021). 10.1038/s41467-021-21246-9

28 Zanet, J. et al. Endogenous Myc controls mammalian epidermal cell size, hyperproliferation, endoreplication and stem cell amplification. J Cell Sci 118, 1693–1704 (2005). 10.1242/jcs.02298

29 Madden, S. K., de Araujo, A. D., Gerhardt, M., Fairlie, D. P. & Mason, J. M. Taking the Myc out of cancer: toward therapeutic strategies to directly inhibit c-Myc. Mol Cancer 20, 3 (2021). 10.1186/s12943-020-01291-6

30 Luo, W. et al. c-Myc inhibits myoblast differentiation and promotes myoblast proliferation and muscle fibre hypertrophy by regulating the expression of its target genes, miRNAs and lincRNAs. Cell Death Differ 26, 426–442 (2019). 10.1038/s41418-018-0129-0

31 Purhonen, J., Klefstrom, J. & Kallijarvi, J. MYC-an emerging player in mitochondrial diseases. Front Cell Dev Biol 11, 1257651 (2023). 10.3389/fcell.2023.1257651

32 Qu, A. et al. Role of Myc in hepatocellular proliferation and hepatocarcinogenesis. J Hepatol 60, 331–338 (2014). 10.1016/j.jhep.2013.09.024

33 Volckaert, T., Campbell, A. & De Langhe, S. c-Myc regulates proliferation and Fgf10 expression in airway smooth muscle after airway epithelial injury in mouse. PLoS One 8, e71426 (2013). 10.1371/journal.pone.0071426

34 Barros, L. et al. CD8(+) tissue-resident memory T-cell development depends on infection-matching regulatory T-cell types. Nat Commun 14, 5579 (2023). 10.1038/s41467-023-41364-w

35 Rubinstein, A. et al. CXCR3-Expressing T Cells in Infections and Autoimmunity. Front Biosci (Landmark Ed) 29, 301 (2024). 10.31083/j.fbl2908301

36 Guo, K. et al. The chemokine receptor CXCR3 promotes CD8(+) T cell-dependent lung pathology during influenza pathogenesis. Sci Adv 10, eadj1120 (2024). 10.1126/sciadv.adj1120

37 Liu, J. et al. Local production of the chemokines CCL5 and CXCL10 attracts CD8+ T lymphocytes into esophageal squamous cell carcinoma. Oncotarget 6, 24978–24989 (2015). 10.18632/oncotarget.4617

38 Groom, J. R. & Luster, A. D. CXCR3 in T cell function. Exp Cell Res 317, 620–631 (2011). 10.1016/j.yexcr.2010.12.017

39 Tokunaga, R. et al. CXCL9, CXCL10, CXCL11/CXCR3 axis for immune activation - A target for novel cancer therapy. Cancer Treat Rev 63, 40–47 (2018). 10.1016/j.ctrv.2017.11.007

40 Cheuk, S. et al. CD49a Expression Defines Tissue-Resident CD8(+) T Cells Poised for Cytotoxic Function in Human Skin. Immunity 46, 287–300 (2017). 10.1016/j.immuni.2017.01.009

41 Johnson, M. D., Witherden, D. A. & Havran, W. L. The Role of Tissue-resident T Cells in Stress Surveillance and Tissue Maintenance. Cells 9 (2020). 10.3390/cells9030686

42 Foote, A. G., Lungova, V. & Thibeault, S. L. Piezo1-expressing vocal fold epithelia modulate remodeling via effects on selfrenewal and cytokeratin differentiation. Cell Mol Life Sci 79, 591 (2022). 10.1007/s00018-022-04622-6

43 Langmead, B. & Salzberg, S. L. Fast gapped-read alignment with Bowtie 2. Nat Methods 9, 357–359 (2012). 10.1038/nmeth.1923

44 Trapnell, C. et al. Transcript assembly and quantification by RNA-Seq reveals unannotated transcripts and isoform switching during cell differentiation. Nat Biotechnol 28, 511–515 (2010). 10.1038/nbt.1621

45 Xie, Z. et al. Gene Set Knowledge Discovery with Enrichr. Curr Protoc 1, e90 (2021). 10.1002/cpz1.90

46 Foote, A. G., Wang, Z., Kendziorski, C. & Thibeault, S. L. Tissue specific human fibroblast differential expression based on RNAsequencing analysis. BMC Genomics 20, 308 (2019). 10.1186/s12864-019-5682-5

47 Satija, R., Farrell, J. A., Gennert, D., Schier, A. F. & Regev, A. Spatial reconstruction of single-cell gene expression data. Nat Biotechnol 33, 495–502 (2015). 10.1038/nbt.3192

48 Becht, E. et al. Dimensionality reduction for visualizing single-cell data using UMAP. Nat Biotechnol (2018). 10.1038/nbt.4314

49 Fan, C. et al. irGSEA: the integration of single-cell rank-based gene set enrichment analysis. Brief Bioinform 25 (2024). 10.1093/bib/bbae243

50 Trapnell, C. et al. The dynamics and regulators of cell fate decisions are revealed by pseudotemporal ordering of single cells. Nat Biotechnol 32, 381–386 (2014). 10.1038/nbt.2859

